# The functional maturation of mouse spermatozoa is underpinned by global remodeling of the cellular phosphoproteome

**DOI:** 10.1101/2025.09.18.677062

**Authors:** David A. Skerrett-Byrne, Amanda L. Anderson, Raffaele Teperino, Nathan D. Burke, Elizabeth G. Bromfield, Matthew D. Dun, Valérie Gailus-Durner, Helmut Fuchs, Susan Marschall, Martin Hrabě de Angelis, Sean J. Humphrey, Brett Nixon

## Abstract

The functional maturation of mammalian spermatozoa is driven by modification of their intrinsic proteome as the cells transit the male (epididymal sperm maturation) and female reproductive tracts (capacitation). Here, high-resolution mass spectrometry was used to interrogate the central role that phosphoproteomic changes play in the functional remodeling of mouse spermatozoa. This strategy identified 14,586 site-specific phosphorylation events, including the phosphorylation of 573 proteins and dephosphorylation of 426 during epididymal maturation and additional phosphorylation changes in 211 proteins linked to capacitation. We identified over 300 kinases that putatively govern these events, including three novel kinases (STK33, HIPK4, and PAK1) implicated in acrosomal exocytosis. The functional relevance of these data was confirmed via the use of knockout mouse models, which demonstrated several phosphoproteins as being essential for sperm motility and fertilization capacity. These findings illustrate that large-scale phosphorylation remodeling occurs during sperm maturation with implications extending to novel means of fertility regulation.

**HIGHLIGHTS:**

- Sperm phosphoproteome is dramatically remodeled during post-testicular maturation
- Identification of >14,000 site-specific phosphorylation events providing comprehensive insight into sperm cell signaling events associated with functional maturation
- Major changes in the sperm phosphoproteome coincide with epididymal maturation whereas capacitation results in more modest changes
- Identification of 343 novel kinases potentially important for conferring functional maturity to spermatozoa and demonstrated role for STK33, HIPK4 and PAK1 kinases
- Knockout mouse models of 23 genes provided *in vivo* validation, with loss of these proteins leading to pronounced defects in sperm motility and fertilization capacity
- All data is available via our interactive ShinySpermPhospho application - https://reproproteomics.shinyapps.io/ShinySpermPhospho/

## INTRODUCTION

Among the most challenging unanswered questions in the field of reproduction are the mechanisms by which spermatozoa acquire the functional competence to fertilize an ovum during their extragonadal development. Indeed, despite enduring a tortuous process of cytodifferentiation within the testes, the highly specialized spermatozoa that are delivered into the male reproductive tract (epididymis) lack the capacity for progressive motility and to recognize an ovum.^1,2^ The potential to fulfil these fundamental roles is instead progressively acquired as the sperm transcend the epididymis and subsequently, the female reproductive tract.^3^ A distinctive hallmark of these sequential phases of maturation is that they occur in the complete absence of *de novo* gene transcription and protein translation and are therefore completely reliant on the modification of the intrinsic sperm proteome.^4–8^ We have recently shown that remodeling of the core sperm proteome encompasses the selective loss and gain of hundreds of proteins.^9^ Notwithstanding the extent of these changes, however, the proteomic landscape of the maturing sperm cell is also exquisitely sensitive to post-translation modifications (PTM). Indeed, of the more than 200 known forms of PTM,^10^ transient protein phosphorylation has emerged as one of the major regulators of sperm function; being implicated as the dominant driving force behind the activation of cellular metabolism, signaling, and surface modifications.^11–15^ Despite this knowledge, the full cascade of phosphorylation events that underpin sperm maturation have yet to be resolved.

To address this challenge, here we have exploited high-resolution tandem mass spectrometry (MS) to characterize proteome-wide changes in the mouse sperm phosphoproteome associated with their transit of the epididymis and elicited in response to capacitation stimuli. This strategy capitalizes on recent advancements in MS-based phosphoproteomic technologies, which are now amenable to the simultaneous high-resolution mapping of multiple site-specific phosphorylation events.^16,17^ Such techniques have been instrumental in resolving the true complexity of this form of PTM and shedding new light on the scale of the biological consequences resulting from dysregulation of protein phosphorylation.^18^ Indeed, it is now apparent that as many as two-thirds of all cellular proteins harbor at least one phosphorylation site. Against this background of daunting complexity, we report the quantification of ∼14,600 site-specific phosphorylation events corresponding to thousands of maturation-related sperm proteins. Specifically, our data reveal largescale phosphorylation and dephosphorylation associated with sperm transit of the epididymis. Thereafter, the capacitation events that are initiated upon contact with the female reproductive tract, elicit relatively modest phospho-changes, suggesting these additional modifications fine-tune sperm function in preparation for fertilization. Moreover, we identified a subset of novel kinases that play key roles in shaping the phospho-status of the maturing sperm cell. Such information provides new insight into the complexity of the signaling processes underpinning sperm transformation into functionally competent cells with important implications for development of novel approaches to male fertility regulation such as non-hormonal contraceptives.

## RESULTS

### Assessment of the key phosphorylated amino acids in maturing mouse sperm

As a prelude to the identification of substrates (de)phosphorylated during the post-testicular maturation of mouse spermatozoa, we first examined the phospho-status of these cells immediately following isolation from the lumen of the caput and cauda epididymis (non-capacitated) and again after *in vitro* incubation under conditions optimized to promote capacitation (Figure 1). Immunoblotting of sperm protein lysates from this experiment confirmed that, unlike that of their functionally mature cauda epididymal sperm counterparts, the immature population of caput epididymal spermatozoa are refractory to capacitation stimuli. Indeed, minimal change in the relative abundance, or overall profile, of phospho-serine, -threonine, - tyrosine or PKA phospho-substrate residues was documented in caput epididymal spermatozoa after incubation in medium supplemented with the capacitation-inducing stimuli, dibutyryl cAMP and pentoxifylline; all of which remained at basal levels comparable to that of freshly isolated spermatozoa (Figure 1). By contrast, pronounced changes in both the phospho-protein profile (Figure 1A-H) and intensity (Figure 1I-P) of antibody labeling were detected in cell lysates prepared from spermatozoa that had completed transit to the distal, cauda, epididymis. Particularly in the case of phosphotyrosine (Figure 1E-F) and phosphoPKA substrates (Figure 1G-H), the intensity of labeling was also notably increased across several prominent proteins spanning the range of ∼20 – 220 kDa upon stimulation of capacitation (arrowheads). More modest responses were noted following phosphothreonine labeling of capacitated cauda epididymal sperm lysates, whereby increased staining appeared largely restricted to bands of ∼100 and 220 kDa (Figure 1C-D; arrowheads). In the case of phosphoserine, virtually all labeled proteins appeared to be constitutively phosphorylated in cauda epididymal spermatozoa such that their labeling intensity did not change under the capacitation conditions employed in this study (Figure 1A-B).

**Figure 1:**
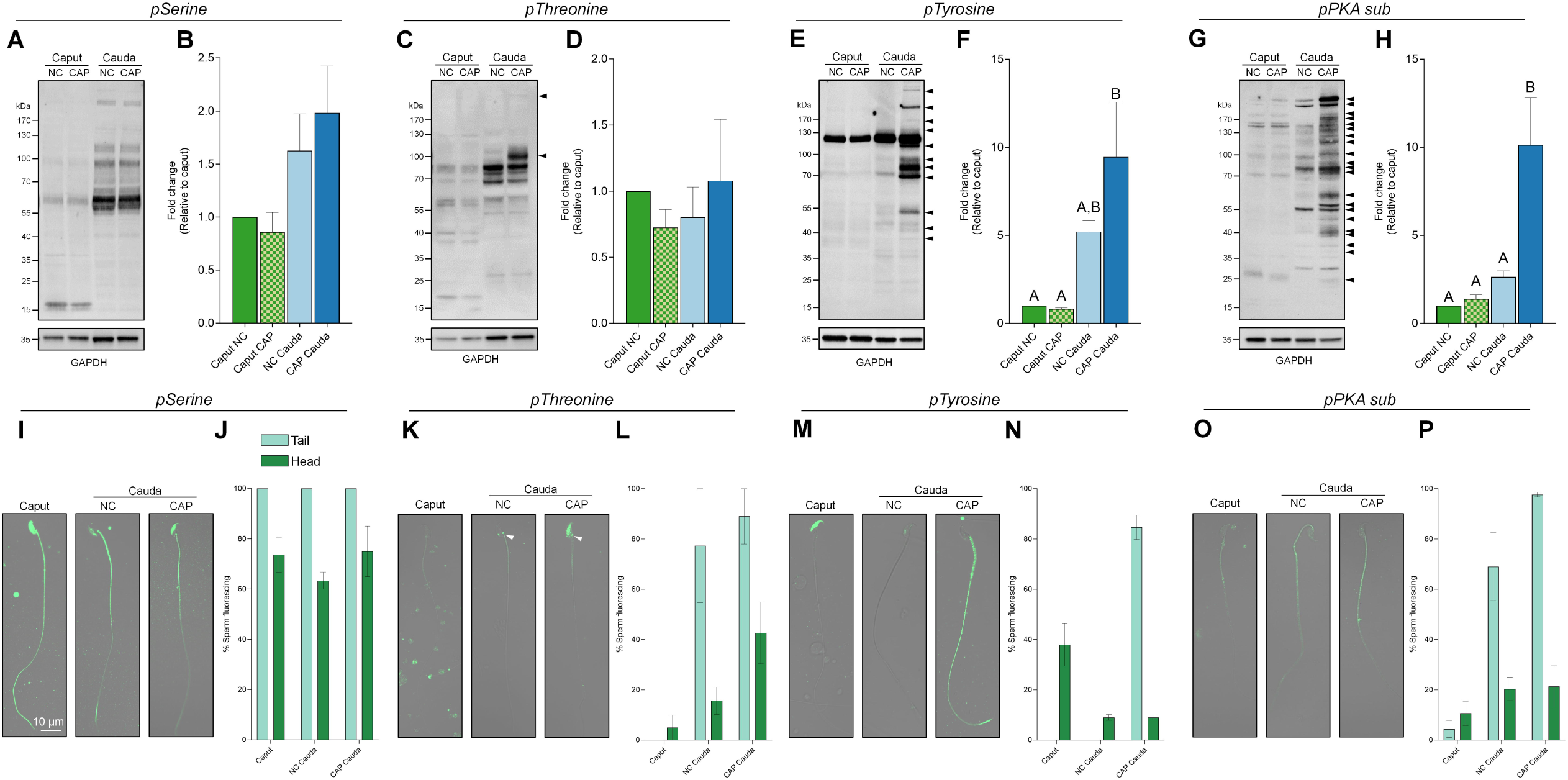
Investigation of phosphorylated status of key amino acids in maturing epididymal mouse spermatozoa. Immature (caput) and mature (cauda) sperm cells were assessed under non-capacitated (NC) or capacitated conditions (CAP) by immunoblot for harboring phosphorylation of the serine (A, E), threonine (B, F), tyrosine (C, G) and substrates of PKA (D, H). Caput and cauda (NC and CAP) sperm cells were further examined by immunocytochemistry with counts taken for fluorescence of the sperm tail and head, looking at phosphorylation of the serine (I, M), threonine (J, N), tyrosine (K, O) and substrates of PKA (L, P); scale bar of 10 mm included. All immunoblotting experiments were repeated with at least three biological replicates. Densitometric data normalization was performed against the loading control protein GAPDH, and each value subsequently expressed as a fold change relative to the caput sperm. Data were analyzed by one way ANOVA with GraphPad Prism.

Complementing these immunoblotting data, immunofluorescent labeling of spermatozoa with anti-phosphoprotein antibodies revealed distinctive patterns of staining dependent upon both the targeted phosphosubstrates and the maturation status of the sperm population. Thus, in the case of phosphoserine, strong flagellum (mid- and principle-piece) and staining was detected in virtually all spermatozoa irrespective of their maturation status (Figure 1I-J). In immature caput epididymal spermatozoa, flagellum labeling was accompanied by additional labeling of the entire sperm head in ∼70% of cells, whereas in the mature cauda epididymal counterparts, head labeling became restricted to the equatorial region; but did not undergo further change as a function of the capacitation status of the cell population (Figure 1I-J). By contrast, both the proportion of phosphothreonine labeled sperm cells and the distribution of labeling was notably influenced by epididymal maturation and subsequent capacitation, such that the negligible staining of caput spermatozoa was replaced with flagellum labeling, albeit of relatively modest intensity, in ∼80% of cauda spermatozoa (Figure 1K-L). Capacitation promoted additional phosphothreonine labeling, particularly in the connecting piece and throughout the head of these mature cells (Figure 1K; arrowheads). Similarly, phosphotyrosine labeling also proved sensitive to sperm maturation such that the equatorial labeling detected in the head of ∼40% of caput spermatozoa was replaced with intense flagellum labeling in >85% of the capacitated cauda sperm population (Figure 1M-N). Finally, the labeling of sperm phospho-PKA substrates increased significantly coincident with epididymal transit and again upon induction of capacitation, with these targets being predominantly localized to the sperm flagellum (mid- and principal-piece) (Figure 1O-P). Importantly, these data proved consistent across each biological replicate assessed.

### Mass-spectrometry based characterization of the maturing mouse sperm phosphoproteome

To expand our analysis to the level of phosphorylation driven changes associated with the functional transformation of mouse spermatozoa, we subjected populations of caput (immature) and cauda (mature) epididymal sperm to high-resolution MS-based phosphoproteomics ^19,20^ (Figure 2A,B). The selected workflow (EasyPhos) involves protein extraction, trypsinization and phosphopeptide enrichment in a 96-well format. The resultant phosphopeptide populations were analyzed using label-free nano liquid chromatography-tandem mass spectrometry (nLC-MS/MS) (Figure 2C). These combined platforms returned an unprecedented depth of coverage for each sperm population with 12,415, 8,240, and 8,156 individual phosphosites (i.e. specific amino acid residues) identified in caput, non-capacitated (NC) and capacitated (CAP) cauda sperm samples, respectively (Figure 2D). These complex inventories mapped to 7,835 (2,229 proteins), 5,166 (1,377 proteins) and 5,144 (1,343 proteins) phosphopeptides in caput, NC and CAP cauda sperm samples, respectively (Table S1-3). Together, a total of 14,586 unique phosphosites were identified in maturing mouse sperm, mapping to 2,496 parent proteins (Figure 2D).

**Figure 2:**
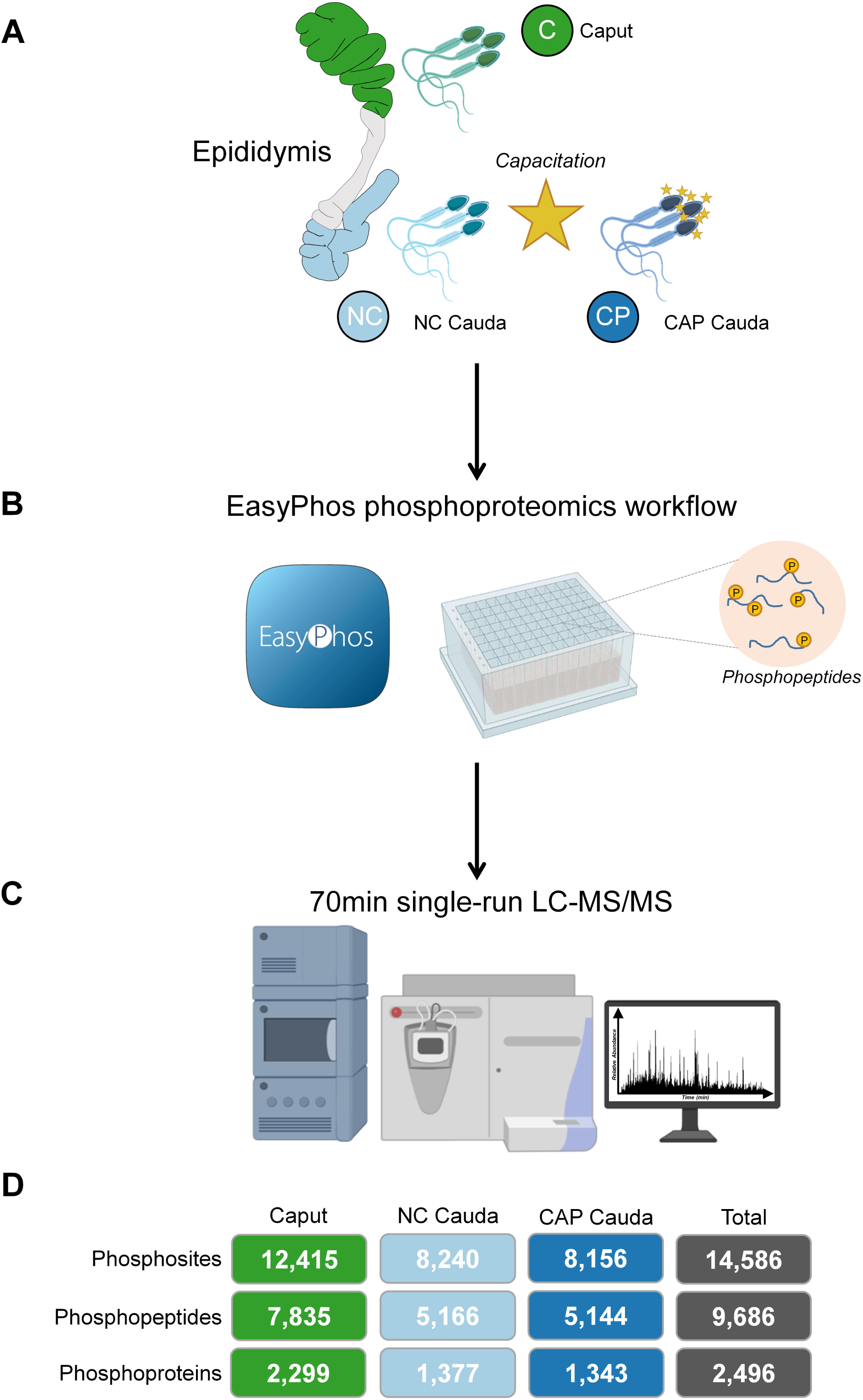
Experimental workflow for phosphoproteomic assessment of functionally immature and mature mouse spermatozoa. (A) Epididymal mouse sperm cells were collected from the proximal caput (green) and distal cauda (light blue), with a proportion of the latter induced to capacitate *in vitro* (dark blue). (B) Each population was subjected to the EasyPhos platform ^20^ to enrich for phosphopeptides before (C) 70 min single run analysis on an Orbitrap HF-X mass spectrometer. (D) Summary of the number of phosphosites, phosphopeptides and their corresponding parent proteins for each population and the sum total of all. Aspects of this figure were created with BioRender.com.

Pearson correlation plots confirmed that the phosphoprotein data arising from each biological replicate were highly reproducible (Figure 3A), with an average correlation of 0.96 across all samples. Additionally, an unbiased principal component analysis (PCA) revealed strong grouping of biological replicates with clear separation between the three different sperm populations analyzed (Figure 3B). Moreover, the PCA plot showcased caput epididymal sperm as being distinct from their cauda epididymal sperm counterparts, with component 1 accounting for 81.2% of this variation. Notably, the cauda epididymal sperm populations were also separated based on their capacitation status. We next characterized the relative proportion of the major phosphorylated residues, namely serine (pS), threonine (pT) and tyrosine (pY), with serine accounting for ∼86% of all phosphorylation sites across each sperm population (Figure 3C). The next most abundant phosphorylated residue was threonine, representing 11.0% of all identified phosphoresidues in caput epididymal sperm and 12.5% in both of the cauda epididymal sperm populations. Lastly, phosphotyrosine initial accounted for 1.3% of all phosphoresidues identified in immature caput epididymal sperm before experiencing a modest increase to 1.6% in mature non-capacitated cauda epididymal sperm and 1.8% (a 38% proportional increase) in their capacitated counterparts. Together, these distributions closely align with 86.4% (pS): 11.8% (pT): 1.8% (pY) ratio commonly observed in somatic cells ^21^. Comparative analysis of the phosphopeptide inventories revealed some 3,675 phosphopeptides (40.2%) were shared across the three sperm populations analyzed, but strikingly, 3,377 phosphopeptides were uniquely identified in caput epididymal sperm (Figure 3D). Cauda epididymal sperm harbored 200 and 267 unique phosphopeptides in non-capacitated and capacitated populations, respectively.

**Figure 3:**
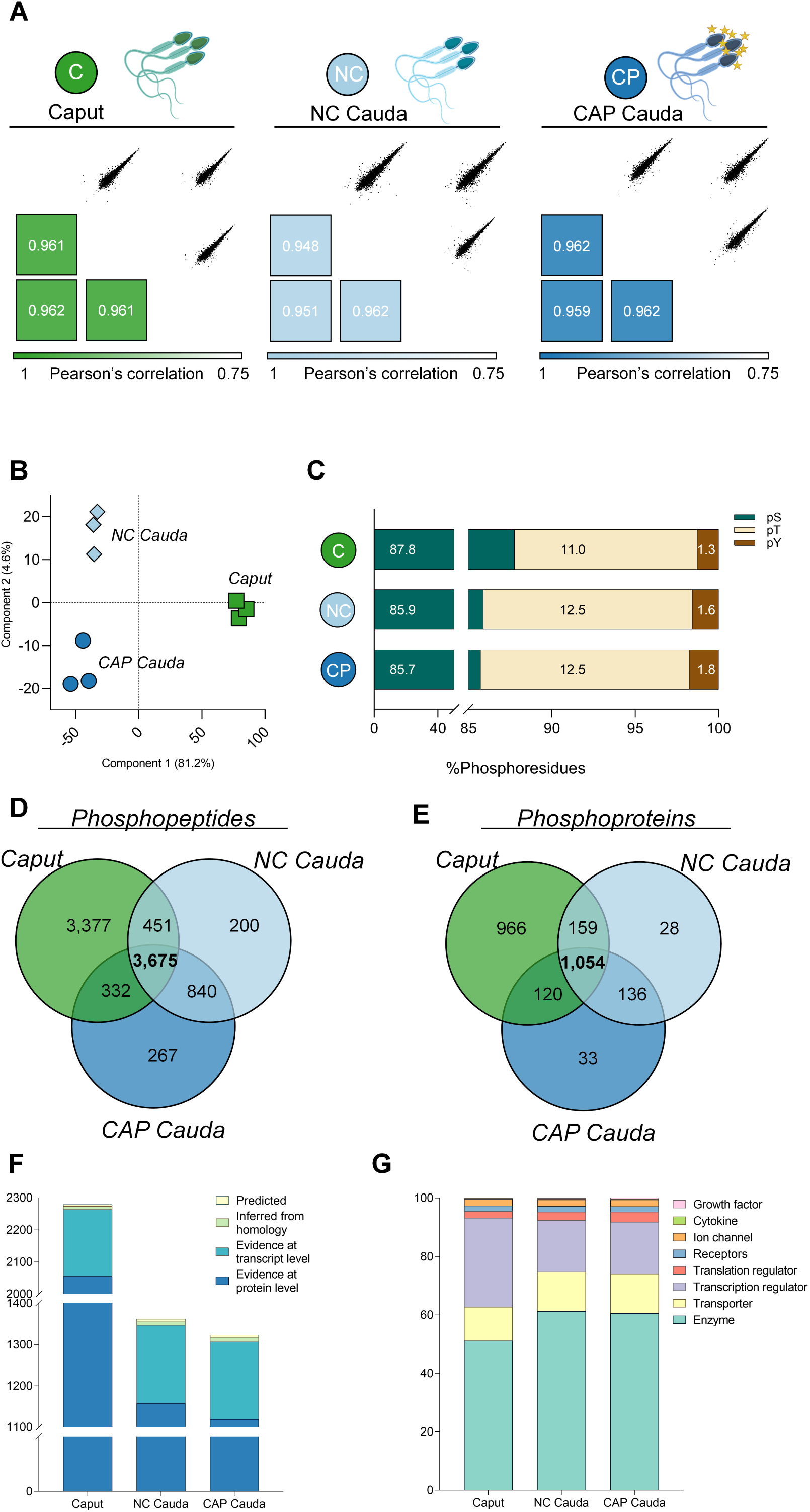
Phosphoproteomic characterization of maturing epididymal sperm populations. (A) Robustness of phosphoproteomic analysis assessed by Pearson correlation plots for caput (immature, green), non-capacitated cauda (mature, light blue), and capacitated cauda (dark blue) epididymal sperm populations. (B) Principal component analysis (PCA) of the full quantitative phosphoproteomic profile of each biological replicate across all three sperm cell population; caput (green squares), NC cauda (light blue diamonds), and CAP cauda (dark blue circles). (C) Proportional bar chart of the distribution of phosphorylated serine (pS), threonine (pT) and tyrosine (pY) detected in each sperm cell population. Venn Diagram of the detected (D) phosphopeptides and their parent (E) phosphoproteins across the three epididymal sperm populations, illustration shared and unique to each. (F) UniProt Knowledge Base annotation for evidence level of protein existence and (G) protein type classification determined by Ingenuity Pathway analysis (IPA).

Similar results were observed at the level of the parent proteins with 42.2% of all phosphoproteins shared between each shared population (Figure 3E). Comparison of these phosphoproteomic data to that arising from a previous proteomic characterization of matching epididymal sperm populations ^19^ revealed a 79.6% overlap, with an additional 510 proteins being uniquely identified in this phosphoproteomic analysis (Figure S1).

Interrogation of the total phosphoproteomic data based on curated evidence supporting the existence of each protein using UniProt annotation ^22^ demonstrated that 2,238 (89.7%) of the identified phosphoproteins have previously been annotated at the protein level, yet a subset of 258 proteins had yet to be associated with evidence at the protein level (Figure 3F). Among this latter group of proteins, the majority (i.e., 240 proteins; 9.6%) have been documented at the transcript level, while the remainder have been either inferred from homology (12 proteins; 0.5%) or predicted to exist (6 proteins; 0.2%) (Figure 3F). Moreover, UniProt annotation revealed 77.2% of the epididymal sperm phosphoproteins (1,928) identified herein have previously been shown to harbor phosphorylation at the residue level. An additional dividend of the resources provided by UniProt coupled with EMBL-EBI GO annotations is the ability to assign cellular location of identified proteins. Accordingly, 64.1% of the phosphoproteins identified herein were able to be putatively mapped to key sperm domains, with the nucleus (1,242 proteins) and mitochondria (149 proteins) accounting for the two largest annotated domains (Figure S2 and Table S1-3). Extending the *in-silico* analyses of identified phosphoproteins, Ingenuity Pathway Analysis (IPA) was used to classify the phosphoproteins according to their function. This strategy identified enzyme as the leading protein classification, accounting for 51.2% of all phosphoproteins identified in immature caput epididymal sperm, and 61.3% and 60.5% in mature NC and CAP caudal epididymal sperm populations, respectively (Figure 3F and Tables S1-3). Notably, the kinases and phosphatases that reciprocally regulate phosphorylation, accounted for 16.1% of the enzymes identified in caput epididymal sperm yet increased following epididymal maturation to comprise 19.8% of all enzymes in both cauda epididymal sperm populations. The majority of the remaining protein classifications remained relatively constant throughout epididymal sperm maturation with the notable exception of transcription regulators, which decreased proportionally by 58% from immature caput sperm (30.5%) to mature cauda sperm populations (17.7%) (Figure 3G and Tables S1-3).

### Epididymal maturation accounts for the majority of maturation-associated sperm cell signaling

To assess temporal changes in the sperm phosphoproteome associated with their functional maturation, statistical analyses (i.e., ANOVA testing) were used to identify differentially phosphorylated peptides across each of the three subpopulations of spermatozoa (i.e., caput and NC and CAP cauda epididymal spermatozoa). This approach revealed 2,975 phosphorylated peptides that were significantly altered in terms of their relative abundance across the three sperm populations (Figure 4A, Table S4; q-value ≤ 0.05). Unbiased hierarchical clustering of these significant phosphorylated peptides identified four distinct groupings of peptides with similar patterns of expression, which we will refer to as Groups from this point on (Figure 4A), namely; Group 1) phosphopeptides that were dephosphorylated during epididymal maturation and remained unchanged during capacitation; Group 2) peptides that were phosphorylated during epididymal maturation and remained unchanged during capacitation; Group 3) phosphopeptides whose phosphorylation status was not changed during epididymal transit but experienced dephosphorylated during capacitation, and finally those; Group 4) peptides whose phosphorylation status was unchanged during epididymal transit but thereafter became phosphorylated during capacitation. Notably, the majority of these phosphorylation changes (86.1%) were established during sperm transit of the epididymis as opposed to during capacitation, such that Groups 1 and 2 featured 743 and 1,818 phosphopeptides, respectively (Table S4); representing a five-fold higher response than that observed during capacitation (Figure 5B). These phosphopeptides mapped to 426 parent proteins in Group 1 and 573 in Group 2, representing 1.7 and 3.2 phosphorylation events per protein, respectively. Following epididymal maturation, capacitation driven changes accounted for the remaining 13.9% of differential phosphorylation (Figure 4A) featured in groups 3 (61 phosphopeptides from 42 parent proteins) and 4 (353 phosphopeptides from 188 parent proteins) (Table S4). Furthermore, several kinases and phosphatases, the key regulators of this important PTM, were identified in each group; Group 1, 20 kinases and 4 phosphatases; Group 2, 40 kinases and 19 phosphatases; Group 3, 1 phosphatase; Group 4, 11 kinases and 10 phosphatases. When compared to the known kinases (Figure 4B) and phosphatases (Figure 4C) present in epididymal sperm ^19^, 91.5% and 89.3% respectively were detected compared to the ANOVA significant phosphopeptides. Five unique kinases and three phosphatases were detected in the phosphoproteome data, including the tyrosine kinases LYN and hepatocyte growth factor receptor (MET) and serine/threonine-protein phosphatase PP1-gamma catalytic subunit (PPP1CC) (Table S4).

**Figure 4:**
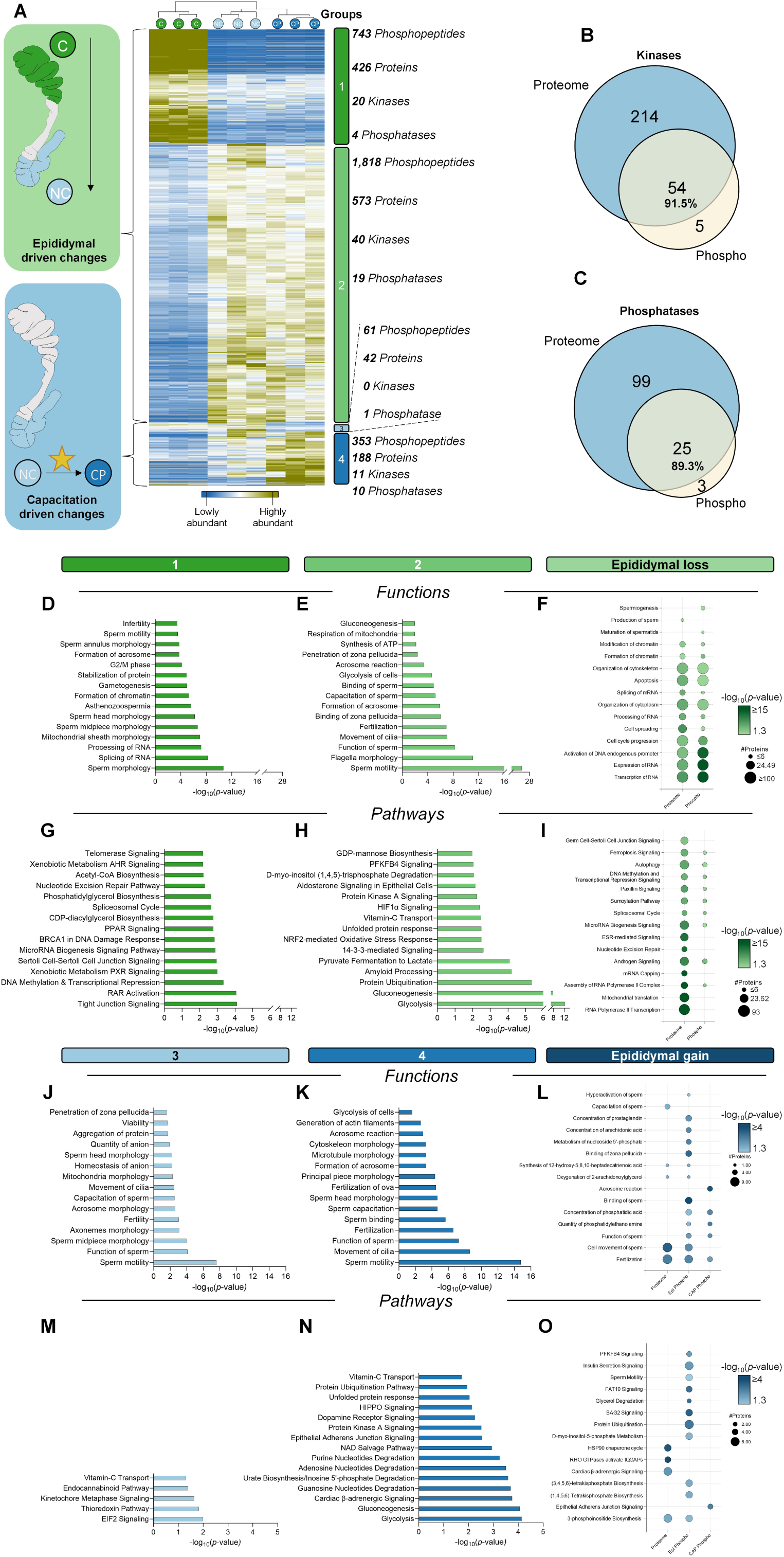
Epididymal transit defines major priming of phosphorylated proteins prior to fine-tuning by capacitation. (A) Hierarchical clustering of ANOVA significant (one-way, *p*-value ≤ 0.05) phosphopeptides from caput (immature, green), non-capacitated cauda (mature, light blue), and capacitated cauda (dark blue) epididymal sperm populations. Groups of significant change related to epididymal maturation are signposted by green boxes 1 & 2, whilst capacitation driven changes are noted by blue boxes 3 & 4. On the right-hand side, are the corresponding numbers of phosphopeptides, their corresponding parent proteins and the number of events/proteins. Each group was individually analyzed by IPA, followed by a comparative analysis of the parent parents. The top 10 significant (*p*-value ≤ 0.05) (B) sperm related molecular functions and (C) pathways area represented as heatmaps.

**Figure 5:**
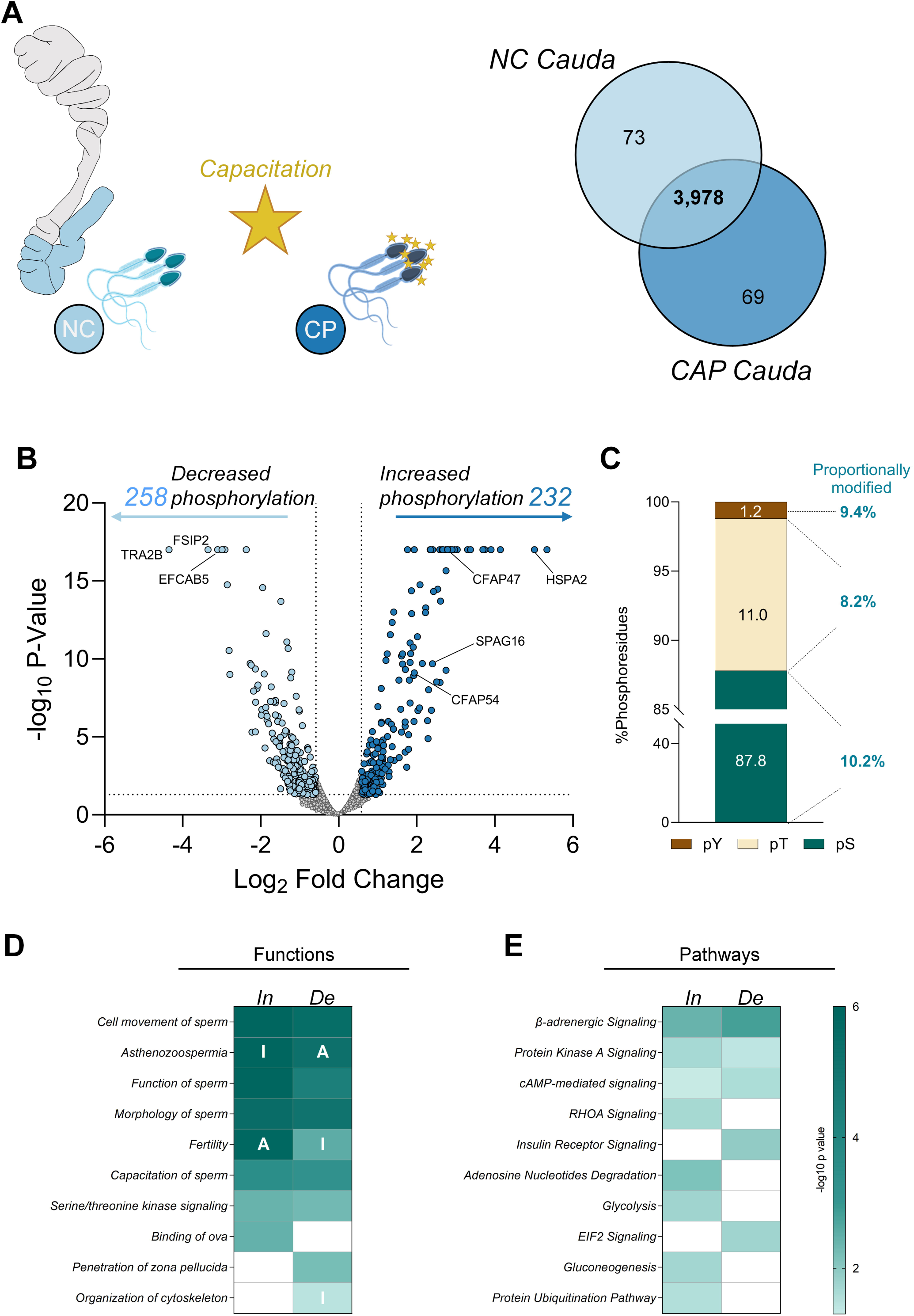
Capacitation elicits sustainable shifts in phosphorylation. (A) Venn Diagram comparison of the phosphopeptides detected within the non-capacitated cauda (mature, light blue), and capacitated cauda (dark blue) epididymal sperm populations. (B) Volcano plot analysis depicting the differential phosphorylation status of 490 phosphopeptides associated with capacitation; log_2_ fold change (x axis) and –log *p*-value (y axis) with thresholds of ± 1.5 fold change and *p*-value ≤ 0.05. (C) Percentage breakdown of the significantly phosphorylated amino acids (serine, threonine and tyrosine), with annotation of the proportional change of each. A comparative functional analysis of the parent parents that experienced significantly increased or decreased phosphorylation following capacitation was carried with IPA. Heatmaps depict the top 10 significant (*p*-value ≤ 0.05) (D) sperm related molecular functions and (E) pathways; A = predicted activation; I = predicted inhibition.

To gain insight into the potential biological impact of temporal changes in sperm phosphorylation status, we interrogated each of the four hierarchical groups using IPA’s phosphorylation knowledge base, focusing our analysis on those pathways linked to male reproduction and sperm function (Figure 4D-O). In accordance with the functional importance of the epididymis in preparing sperm for fertilization, Groups 1 and 2 were characterized by a dephosphorylation of proteins with functions related to the legacy of testicular maturation (Figure 4D) and the reciprocal phosphorylation of proteins critical for sperm related functional categories (Figure 4E). By way of illustration, the most enriched (*p*-value ≤ 0.05) functional categories in Group 1 (i.e., those peptides dephosphorylated during epididymal maturation) included those associated with morphological specialization (e.g. ‘sperm midpiece’ and ‘sperm head’) and several RNA related functions (e.g. ‘splicing of RNA’ and ‘processing of RNA’) (Figure 4D). In contrast, the enrichment of functional categories in Group 2 (i.e. featuring >570 proteins whose phosphorylation increased during epididymal maturation) included those required for ensuring fertilization competence (e.g. ‘sperm motility’, ‘function of sperm’, ‘binding of zona pellucida’, and ‘binding of sperm’) (Figure 4E). Given our previous findings of significant protein loss during epididymal maturation ^19^, the subset of proteins that experienced reduced phosphorylation (Figure 3 D-E) were compared to those proteins known to shed along epididymal maturation (Figure 4F). In congruence with proteomic changes, proteins with reduced phosphorylation mapped heavily to functions more relevant to testicular maturation, namely ‘transcription’, ‘expression’ and ‘splicing of RNA’, as well as ‘formation’ and ‘modification of chromatin’ (Figure 4F).

Expanding this *in silico* analysis to focus on cellular pathways revealed that proteins mapping to Group 1 were substantially enriched for upstream testicular processes including ‘Sertoli cell-Sertoli cell junction’ and ‘tight junctions’, and those that shape the sperm epigenome such as ‘DNA methylation and transcriptional repression’ and ‘microRNA biogenesis signaling’ (Figure 4 G, I).^23^ Conversely, those proteins within Group 2 whose phosphorylation was stimulated within the epididymis were associated with cellular pathways crucial for sperm motility, capacitation, and acrosome reaction, specifically ‘PKA signaling’, and ‘protein ubiquitination’. Moreover, in recognition that the cauda epididymis is the primary site for sperm storage, several of these pathways were also linked to the maintenance of cell quiescence and survival, including metabolic pathways such as ‘glycolysis’ and ‘gluconeogenesis’ and antioxidant defense pathways such as the ‘NRF2-mediated oxidative stress response’ (Figure 4H).

Subsequent to epididymal maturation, capacitation was responsible for ∼14% of the recorded changes in protein phosphorylation status that reached statistical significance (Figure 4A). Interrogation of this cohort of proteins revealed a reciprocal enrichment of sperm functions between Groups 3 (dephosphorylated during capitation) and 4 (phosphorylated during capitation), suggesting careful coordination of opposing phosphorylation events are necessary to ensure successful ‘motility’, ‘capacitation’, and ‘acrosome reaction’ (Figure 4J-K). Unsurprisingly, these functions were dominated by phosphorylation (Group 4), involving unique enrichment of proteins involved in ‘fertilization’, ‘glycolysis’, and ‘sperm binding’ (Figure 4K). A comparison of proteins apparently gained during epididymal transit ^19^ with those that are phosphorylated exclusively during epididymal transit and capacitation (Figure 4L) revealed the enrichment of ‘fertilization’, ‘cell movement of sperm’, ‘hyperactivation’ and ‘binding of zona pellucida’ categories, each of which are fundamentally important for the attainment of functional competence. Moreover, a cohort of proteins implicated in the ‘acrosome reaction’ were found to be exclusively phosphorylated in the capacitated sperm population.

Due to the comparatively smaller size of Group 3 (42 proteins), only five pathways (including ‘EIF2 signaling’ and ‘vitamin-C transport’) were significantly enriched in this cohort of proteins that were dephosphorylated during capacitation (Figure 4M). Conversely, Group 4 proteins, which were phosphorylated in response to capacitation, mapped to a rich complement of metabolic pathways (‘glycolysis’ and ‘gluconeogenesis’) and those involved in protein folding / cytoskeletal dynamics (‘unfolded protein response’, ‘protein ubiquitination’ and ‘HIPPO signaling’) (Figure 4N). Returning to the integrative comparison between proteins uniquely gained during epididymal maturation and subsequently phosphorylated during capacitation (Figure 4O), revealed an enrichment of pathways associated with ‘HSP90 chaperone’ and ‘activation of IQGAP’ (IQ motif-containing GTPase-activating proteins), the latter of which is important in actin cytoskeleton dynamics. Similarly, this cohort of proteins were also associated with energy metabolism in nutrient limited environments (‘glycerol degradation’), regulating intracellular calcium levels (inositol phosphate metabolism) and cellular stress response (‘BAG2’), as well as protein ubiquitination (‘FAT10’) (Figure 4M).

### Capacitation-associated sperm protein phosphorylation

Consistent with the translationally inert status of epididymal spermatozoa, previous studies have shown that capacitation leads to only subtle alterations in the overall sperm proteome ^19^. Such findings underscore the potential importance of PTMs such phosphorylation events in terms of facilitating the final priming of spermatozoa in preparation for fertilization. Despite this, our initial comparison of phosphopeptides detected in populations of non-capacitated and capacitated cauda epididymal sperm revealed a high degree of conservation (3,978 phosphopeptides, 96.6%) with only a relatively modest 73 phosphopeptides being lost and a further 69 phosphopeptides gained as a consequence of capacitation (Figure 5A, Table S5). Examination of the conserved phosphopeptides demonstrated that 258 were dephosphorylated following capacitation [fold-change (FC) ± 1.5, *p*-value ≤ 0.05], with the largest changes associated with phosphopeptides from transformer-2 protein homolog beta (TRA2B; FC = –20.4), fibrous sheath-interacting protein 2 (FSIP2; FC = –10.2) and EF-hand calcium-binding domain-containing protein 5 (EFCAB5; FC = –10.2) (Figure 5B). Complementing these dephosphorylation events, we observed increased phosphorylation of 232 peptides following sperm capacitation. As anticipated, these latter phosphopeptides mapped to proteins with known roles in capacitation, including heat shock-related 70 kDa protein 2 (HSPA2; FC = 32.4; implicated in sperm surface remodeling^24,25^) and sperm motility, including sperm-associated antigen 16 protein (SPAG16; FC = 5.3) and several cilia and flagella-associated proteins (CFAP) 44 (FC = 3.1), 47 (FC = 7.9), 57 (FC = 3.6), 100 (FC = 3.5) and 119 (FC = 2.0). Notably, the distribution of phosphorylated amino acid residues that were significantly altered as a consequence of capacitation was retained at the expected ratio of 87.8 serine: 11.0 threonine: 1.2 tyrosine (Figure 5C and Figure 3C). However, analysis of the proportional increase in the phosphorylation of each residue revealed relatively consistent changes with 10.2% of serine residues, 8.2% of threonine residues and 9.4% of tyrosine residues increasing in their phosphorylation status coincident with capacitation (Figure 5C).

To elucidate the biological implications of capacitation-associated changes in phosphorylation, we interrogated the parent proteins of the significantly phosphorylated and de-phosphorylated peptides independently using IPA, as previously described.^26^ This was, in part, due to the limitation of IPA to assess multiple phosphorylated proteins at once, whereby it defaults to the maximum change observed for the parent protein. This analysis included proteins uniquely phosphorylated in either NC or CAP cells. Initial functional analysis revealed, unsurprisingly, a predominance of reproductive functions, including ‘cell movement of sperm’, ‘function of sperm’ and capacitation of sperm’ (Figure 5D). Despite fewer proteins associated with increased phosphorylation, each function was more enriched (i.e., greater *p*-value) in this population than the dephosphorylated changes, e.g. ‘cell movement of sperm’ *p*-value = 4.57 x 10^-13^ and 3.2 x 10^-6^, respectively. Notwithstanding this enrichment, the fact that the pool of phosphorylated proteins recovered from both non-capacitated and capacitated populations of spermatozoa mapped to overlapping functional categories highlights the precise cellular regulation of capacitation, with the reciprocal action of kinases / phosphatases likely holding the key to fine tuning the sperm function in preparation for fertilization. Illustrative of this, we observed ‘binding of ova’ to be exclusively mapped to peptides that experienced increased capacitation-associated phosphorylation whereas ‘penetration of zona pellucida’ was mapped to peptides that were dephosphorylated during capacitation.

Additional pathway analysis demonstrated that a subset of hyperphosphorylated proteins (defined as those with significantly increase phosphorylation, 24 phosphosites) were implicated in pathways crucial for the regulation of acrosomal reaction (i.e., ‘RHOA signaling’ ^19^) and energy metabolism (i.e., ‘glycolysis’ and ‘gluconeogenesis’) (Figure 5E). In contrast, dephosphorylated proteins were uniquely associated with ‘insulin receptor signaling’ and ‘Eukaryotic Initiation Factor 2 (EIF2) signaling’. Shared pathways were those critical for sperm energy metabolism, including ‘Protein Kinase A signaling’ and ‘cAMP-mediated signaling’ (Figure 5E).

### Novel sperm kinase regulators of the acrosome reaction

To begin to leverage the reported phosphoprotein data to deliver biological insight into the control of sperm function, we initially focused on identification of the kinases responsible for promoting upregulation of protein phosphorylation owing to their potential as druggable targets for fertility regulation. Using complementary *in silico* analyses ^26^, including IPA, PhosphoSitePlus,^27^ in conjunction with published epididymal sperm proteome data,^19^ we identified 343 kinases mapping to Groups 2 and 4, as putatively contributing to the increased phosphorylation associated with sperm maturation (Figure S3,Table S6). Given the challenges of characterizing the distinctive role of several kinases in the context of epididymal maturation, we elected to focus on a more tractable approach by analyzing those kinases associated with capacitation (i.e., kinases implicated in the increased phosphorylation of proteins mapping Group 4, each of which displayed increased phosphorylation in response to capacitation stimuli (Figure 4A, Table S6)). This included 25 kinases spanning six major kinase families, including the dominant calcium/calmodulin-dependent protein kinases (CAMK) and AGC families, which together accounted for 48% (12 kinases) of the capacitation-associated kinases (Figure 6A). Notably, this subset of kinases also included several known regulators of sperm function including Protein Kinase A (PKA), Protein Kinase C (PKC), and RAC-alpha serine/threonine-protein kinase (AKT), thus adding credence to the accuracy of this analysis. To further validate this approach, we elected to focus on the functional role of three kinases with hitherto unknown roles in the regulation of sperm capacitation, namely: serine/threonine-protein kinase 33 (STK33), homeodomain-interacting protein kinase 4 (HIPK4) and serine/threonine-protein kinase PAK 1 (PAK1) (Figure 6A).

**Figure 6:**
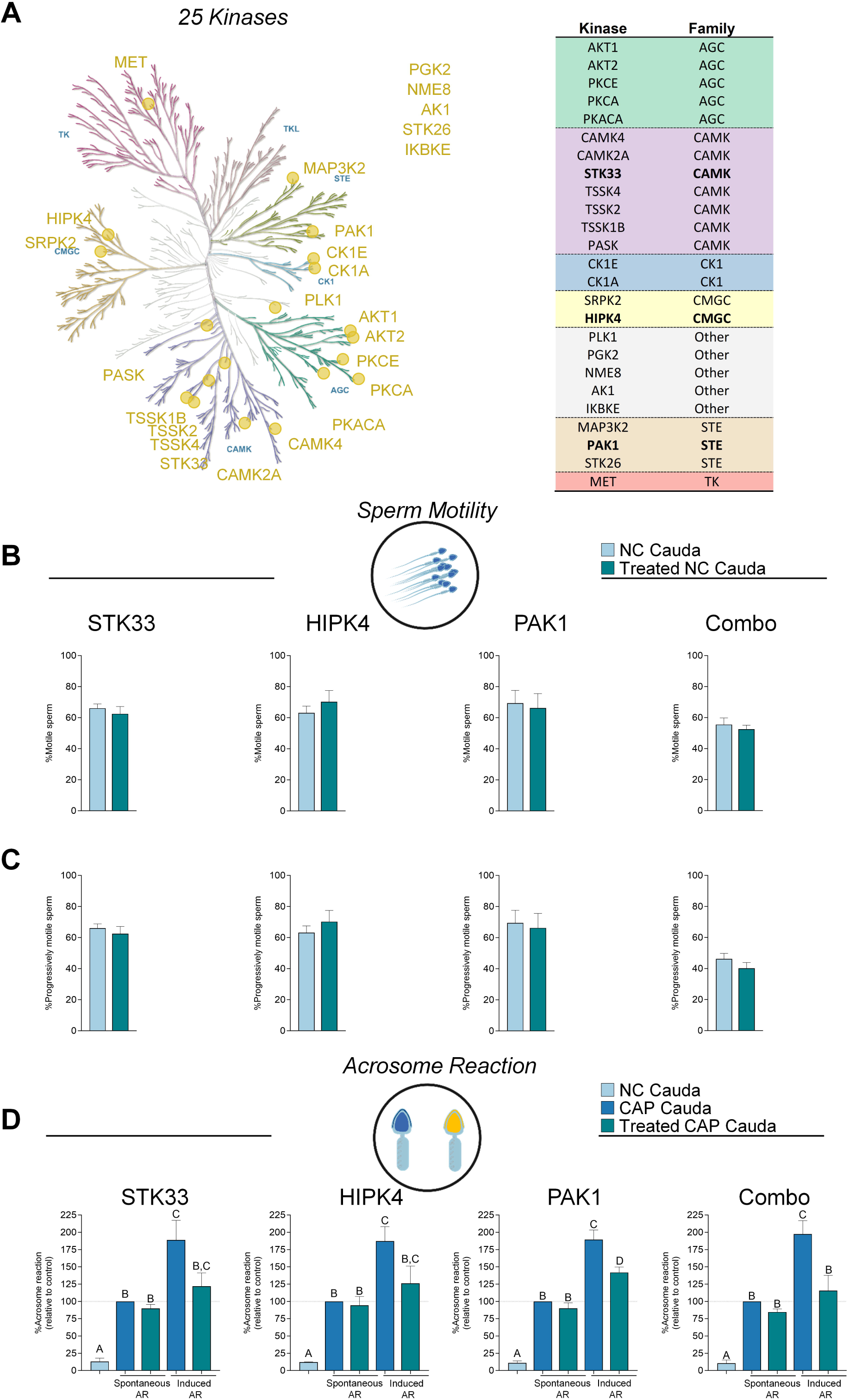
Selective pharmacological inhibition reveals specific functional roles for STK33, HIPK4 and PAK1 in sperm competency. (A) Capacitation driven hyperphosphorylation group 4 (from Figure 5A) was subjected to upstream kinase analyses and a total of 25 kinases were identified and highlight by gold circles in the kinome (KinMap)^80^ with supporting table. (B) Mature non-capacitated cauda spermatozoa (NC) were subject to overall motility and (C) progressive motility counts (expressed as percentage) following 1 h with or without each individual inhibitor or combo; STK33 (ML281), HIPK4 (Foretinib) and PAK1 (NVS-PAK1-1). (D) Acrosome reaction was tested in both spontaneous conditions and induced (A23187 calcium ionophore), in NC and capacitated (CAP) cauda spermatozoa populations, with the latter tested additionally with ML281, Foretinib and PAK1.

To determine the contribution of each target kinase to the regulation of capacitation, we subjected cauda epididymal spermatozoa to selective pharmacological inhibition of STK33 (ML281), HIPK4 (Foretinib) and PAK1 (NVS-PAK1-1) and subsequently assessed their individual and combinatorial effect on sperm motility parameters, ability to complete an acrosome reaction and engage in zona pellucida binding (Figure 6B-D). Prior to undertaking functional analyses, the cytotoxicity of each inhibitor was tested, whereby mouse spermatozoa were incubated for up to 1 h with a range of concentrations for ML281 (14 nM – 14 µM; IC_50_ = 14 nM), Foretinib (0.4 nM – 400 nM; IC_50_ = 0.4 nM), and NVS-PAK1-1 (5 nM - 5 µM; IC_50_ = 5 nM). Importantly, none of the selected kinase inhibitors elicited a significant adverse effect on mouse sperm viability (Figure S4) or motility (Figure S5). Indeed, even at the highest concentrations tested, we observed no significant change in total or progressive sperm motility irrespective of whether the inhibitors were applied individually or as a cocktail (Figure 6B-C). By contrast, the introduction of each inhibitor to populations of capacitating spermatozoa led to a significant reduction in the ability of these cells to complete a calcium ionophore (A23187) induced acrosome reaction; STK33 = 67.1% reduction, *p*-value 0.057; HIPK4 = 60.8% reduction, *p*-value 0.091; PAK1 = 43.7% reduction, *p*-value 0.006 (Figure 6D). The greatest effect was witnessed when spermatozoa were exposed to a cocktail of the three inhibitors; a treatment that effectively returned the rate of acrosomal exocytosis to basal levels equivalent to those witnessed in non-capacitated sperm controls (83.0% reduction, *p*-value 0.005) (Figure 6D). Notably, these responses occurred without an attendant impact on rates of spontaneous acrosome reaction. Collectively these data demonstrate the utility of the phosphoproteomic analytical strategy reported herein in terms of identifying the contribution of novel cellular signaling pathways, and the kinases that regulate them, toward the acquisition of sperm function during their post-testicular development.

### Knockout mouse models confirm important roles for the identified phosphoproteins in sperm function

To further support the importance of proteins whose phosphorylation status was changed during sperm maturation, we accessed the European Mouse Mutant Archive (EMMA) ^28^ to retrieve phenotypic data on sperm motility and *in vitro* fertilization (IVF) resulting from knockout (KO) of 23 proteins of these 47 phospho-targets (Figure 7, Table S7). Importantly, aconitate hydratase mitochondrial (*Aco2*), protein ELYS (*Ahctf1*), UMP-CMP kinase (*Cmpk1*), Rab GDP dissociation inhibitor beta (*Gdi2*), nucleoporin NUP35 (*Nup35*), and protein polybromo-1 (*Pbrm1*) were found to be homozygous lethal, thus limiting our assessment to heterozygous males. Focusing first on sperm motility, we noted that KO of coiled-coil domain-containing protein 183 (*Ccdc183*), glycogen synthase kinase-3 alpha (*Gsk3a*) and ATP-dependent RNA helicase A (*Dhx9*) all led to proportional reductions in total motility of 32.5%, 19.3% and 14.5% respectively, compared to that of sperm from wildtype males (Figure 7A). Such changes were even more pronounced in the context of forward progressive motility, with five KO lines displaying proportional reductions of >28% in this parameter, including *Ccdc183*, *Aco2*, *Gdi2*, *Gsk3a* and optineurin (*Optn*); noting that spermatozoa from the *Optn* KO males exhibiting the most severe phenotype equating to a 40.4% reduction (Figure 7A).

**Figure 7:**
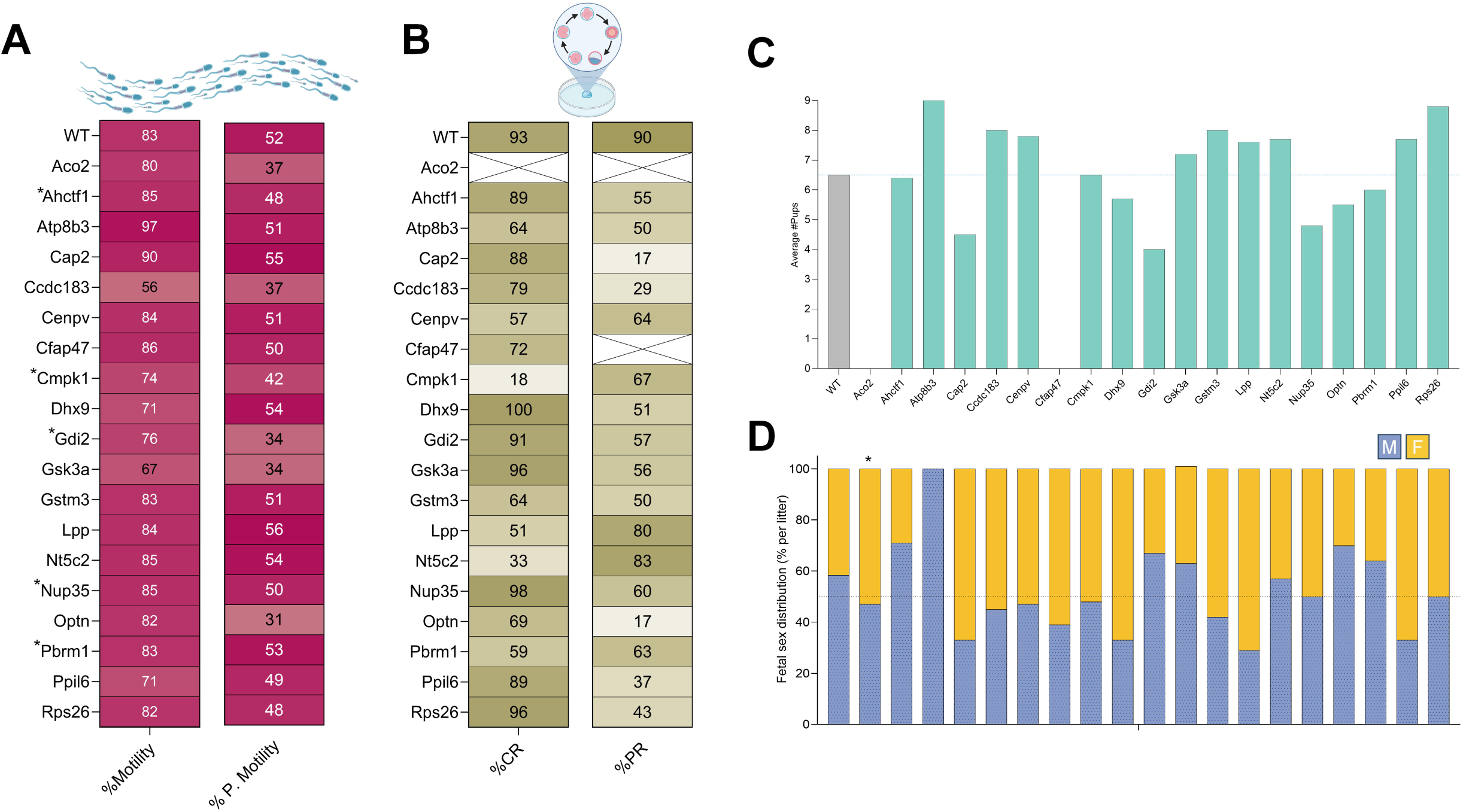
Knock-out mouse lines reveal affects to sperm motility and fertilization capacity. Gene knockout (KO) models were generated and sperm from homozygous males were used for *in-vitro* fertilization (IVF), in total 19 KO lines and wildtype (WT) (Full list of genes is available in Table S7). From these IVF experiments, the heatmaps represent the percentage of (A) motile sperm and those with progressive motility for each KO compared to WT. (B) Fertilization capacity was tracked and heatmaps depict the 2-cell stage cleavage rate (%CR), and pregnancy rate (%PR). Following successful births, the (C) litter size and (D) fetal sex distribution was recorded. * heterozygous males were used instead, as homozygous embryonical lethal or sperm could not fertilize an egg

In seeking to extend the analysis of the fertilization capacity of spermatozoa from KO models we next evaluated their ability to achieve *in-vitro* fertilization (IVF). An initial focus on the cleavage rate to 2-cell embryo stage revealed that nine KO lines experienced marked reductions of >29% compared to fertilization with WT sperm (Figure 7B). Notably, spermatozoa from *Aco2* KO males were rendered incapable of fertilization, whilst *Cmpk1* (80.6%), Cytosolic purine 5’-nucleotidase (Nt5c2; 65.5%) and lipoma-preferred partner homolog (*Lpp*) (45.2%) exhibited the largest decline in cleavage rate (Figure 7B). Following embryo transfer, pregnancy rates were diminished across all KO lines relative to WT, ranging from modest reductions in *Odad3* (85% success) to complete absence of pregnancies in cilia and flagella-associated protein 47 (*Cfap47*) KOs (Figure 7B). Notably, embryos generated from the spermatozoa of *Optn* and *Cap2* KO males achieved pregnancy in only 17% of transfers.

Amongst the pregnancies achieved, litter sizes were largely within expected ranges, though with some deviation for phospholipid-transporting ATPase IK (*Atpb8b3*) KOs producing slightly higher average litter sizes of 9, whereas *Gdi2* yielded smaller litters (average of 4) (Figure 7C). Fetal sex distributions also showed skewing in select KO lines. In this context, litters sired by males from *Cap2*, *Dhx9*, *Ppil6* and *Lpp* KOs exhibited a bias towards female homozygous pups (67% in all and 71% in *Lpp*), while conversely, *Gdi2*, *Optn* and *Ahctf1* tended towards more male heterozygous pups (67%, 70% and 71% respectively) (Figure 7D). Strikingly, in *Atp8b3* KO litters, no female homozygous offspring were observed among the 50 offspring assessed, suggesting possible sex-specific lethality or transmission distortion. Collectively, these data further confirm that sperm proteins targeted for phosphorylation during either epididymal sperm maturation and/or capacitation contribute substantially to key facets of sperm function, including motility, fertilization potential, and reproductive success.

## DISCUSSION

Following their release from the testicular microenvironment, the highly differentiated population of spermatozoa must complete an onerous journey of functional maturation before acquiring the competence to engage in the fertilization cascade. Central to this functional transformation is the modification of the innate sperm proteome; a process that is inextricably linked to cellular signaling cascades occurring in the complete absence of *de novo* gene transcription or protein translation.^1,19^ Here, we provide new insight into the complexity of these modifications, revealing the identification of 2,496 sperm proteins that are subject to 14,586 site-specific phosphorylation events. Building upon previous work examining phosphorylation in epididymal spermatozoa,^29^ we report an overlap of 61% at the peptide level (1,368 phosphopeptides), while also adding 5,659 previously unreported phosphopeptides, thus providing deeper insight into modification of the phosphoproteomic landscape linked to the functional maturation of spermatozoa. Notably, our study also differentiates the scale of phosphoproteomic changes associated with both epididymal transit and capacitation phases of post-testicular maturation, revealing that the majority (86.1%) of (de)phosphorylation events accompany sperm passage through the epididymis as opposed to comparatively modest changes elicited in response to capacitation stimuli. Collectively, such findings emphasize the overall importance of the epididymal driven changes in the sperm proteome in terms of endowing the cells with the potential for fertilization. Thereafter, capacitation appears to fine-tune sperm function to prime the cells to navigate the female reproductive tract, recognize and fertilize an ovum.

In what has become a well-established paradigm, the proteomic architecture of mammalian spermatozoa is subject to substantive remodeling during their successive phases of development. Among the most pronounced of these changes occurs as sperm descend through the epididymis, a process that we have recently shown is accompanied by the combined loss and gain of several thousand sperm proteins.^19,30^ Building upon the detailed proteomic blueprint assembled from these previous studies, here we provide evidence of an additional 510 sperm proteins, elevating the total identified inventory of sperm proteins to 6,891; a number that represents ∼75% of the predicted complexity of the mouse and human sperm proteomes.^31^ Moreover, amongst these 510 proteins, 49 have not been previously confirmed at the protein level, indicating they may represent cell-specific proteins.^32^ Traditionally, phosphorylation has been primarily associated with capacitation^33,34^ with minimal changes recorded during epididymal maturation.^35,36^ However, with advancements in label free phosphoproteomic technology ^26,37^ (noting that label based strategies can mask uniquely detected peptides^29^), here we have revealed the complexity of protein phosphorylation that occurs with the attainment of the potential for sperm fertilization competence during epididymal transit. This strategy also enabled us to differentiate the relative contribution of tyrosine phosphorylation, which in nominally viewed as the key driver of capacitation,^11,38^ versus that of a surprisingly complex and dynamic regulation of serine and threonine phosphorylation; changes that have hitherto been largely overlooked as a consequence of using traditional immunoblotting strategies to study sperm maturation.

Consistent with a progressive refinement of the global sperm proteome,^19^ we noted a 43.1% loss/reduction in the number of phosphopeptides detected in mature cauda epididymal spermatozoa versus that of their immature caput epididymal counterparts. Notably, among those phosphosites conserved across all stages of sperm maturation, 86% underwent significant changes in abundance during epididymal transit; a finding that reinforces the notion that epididymal transit is an intensely active period of molecular programming linked to silencing of the machinery needed for testicular development, delivery of functionally competent spermatozoa and the maintenance of these cells in a quiescent state compatible with their prolonged storage in the epididymal duct.^39–44^ Indeed, we identified over 2,300 phosphopeptides that are significantly altered as sperm transit the epididymis (i.e., Groups 1 and 2); representing >5.5 fold more phosphorylation changes than those documented during downstream capacitation. Functional categorization of these phospho-changes confirmed the extensive dephosphorylation of proteins associated with obsolete functions, such as mRNA processing, spliceosome assembly, and chromatin modification.^42,44^ We interpret these data as putative evidence of silencing of redundant molecular machinery in favor of cellular functions crucial for downstream fertilization. In this way, the epididymis actively contributes to preserving sperm viability while simultaneously endowing the cells with the potential to navigate the formidable barriers they must overcome *en route* to fertilization of an ovum.^1,45^ Accordingly, we identified some 573 proteins that experienced increased phosphorylation during epididymal transit. Such proteins mapped to functional categories deemed essential for fertilization, including sperm motility, acrosome reaction, and binding and penetration of zona pellucida; the latter of which was exclusively identified in the context of those sperm proteins which increased phosphorylation during sperm maturation. This is exemplified by IQ and ubiquitin-like domain-containing protein (IQUB), an evolutionarily conserved protein critical for motility.^31,46^ IQUB is detected at equivalent levels in both immature and mature epididymal sperm,^47^ but shown here to be selectively hyperphosphorylated during epididymal transit and capacitation. In addition, we identified a subset of phosphoproteins critical for sperm motility, capacitation, and zona pellucida binding that are uniquely acquired during epididymal maturation, presumably enabling sperm to respond to subsequent capacitation signals effectively.

Beyond IQUB, we identified a suite of fertilization-related proteins that become phosphorylated during epididymal maturation and maintain these modifications through capacitation, consistent with a priming mechanism. Notably, these included subunits of the CatSper channel (CATSPER3/4/B), which mediate Ca² influx essential for hyperactivation,^48^ PKA catalytic subunit (PRKACA), the central driver of capacitation signaling,^49^ and of motility regulators such as Ropporin-1 (ROPN1),^50^ and t-complex protein 11 (TCP11),^51^ which together likely prepares the flagellar machinery for rapid activation. Zona pellucida-binding proteins were also represented, including sperm surface protein Sp17 (SPA17), a well-characterized sperm– zona adhesion molecule,^52^ and subunits of the CCT chaperonin complex (CCT2/3/4/7, TCP1), which assist in protein folding and zona pellucida interaction.^53^ Finally, phosphorylation of PPP3CC, a sperm-specific calcineurin subunit essential for sperm motility and zona penetration,^54,55^ underscores the importance of epididymal priming for ensuring fertilization competence. Collectively, these data highlight an anticipatory strategy in which phosphorylation events during epididymal transit “switch on” critical elements of the fertilization machinery, equipping spermatozoa to rapidly respond to capacitation cues within the female tract.

In agreement with the proteomic profile of maturing epididymal sperm,^19^ the phosphoproteome is selectively streamlined during epididymal maturation, encompassing an apparent silencing of pathways that are no longer essential for the final stages of fertilization.

This targeted silencing presumably eliminates redundant or obsolete cellular machinery related to earlier sperm morphology maturation. For instance, proteins involved in mRNA processing, spliceosome assembly, and interactions with Sertoli cells are reduced or removed, reflecting the transition from morphological developmental requirements to a streamlined fertilization-ready state. The loss of these functions correlates with improved fertilization capacity, as retention of certain proteins beyond the epididymal phase of development is linked to fertility issues,^19,30^ reinforcing the importance of this selective proteomic refinement. This selective silencing of non-essential functions likely minimizes the molecular load within spermatozoa, enabling the cells to instead dedicate energy towards critical fertilization-related processes like motility, capacitation, and the acrosome reaction as highlighted above. The selective loss of proteins and phosphosites may also enhance the efficiency of responses to signals encountered in the female reproductive tract and during fertilization more broadly. By reducing the complexity of the sperm proteome^19^ and phosphoproteome, epididymal maturation may effectively remove pathways that would otherwise divert resources or create unnecessary metabolic burden while also minimizing the burden of cellular stress that could otherwise compromise fertilization potential.. This refinement process highlights the evolutionary sophistication of epididymal maturation in crafting a highly specialized cell capable of fulfilling its primary roles of reaching, recognizing, and interacting with the oocyte.

Beyond preparing sperm for fertilization, epididymal maturation involves a carefully orchestrated stabilization of sperm function compatible with long-term storage.^56,57^ A central challenge therefore is how to prepare sperm to meet the energetic demands of fertilization while simultaneously maintaining their dormancy and preventing premature activation or exhaustion of metabolic substrates within the male reproductive tract. The phosphoproteomic data presented herein provides a compelling molecular explanation as to how this delicate balance may be achieved. Thus, we anticipate that a key contributor to sperm storage is the inhibitory action of protein dephosphorylation, which essentially places the “*brakes*” on sperm metabolism and motility, effectively rendering the cells quiescent compatible with long-term storage. Supporting this model of metabolic suppression, we observed extensive dephosphorylation of glycolytic enzymes during epididymal maturation, including aldolase A (ALDOA), known to catalyze a rate-limiting step in glycolysis,^58^ phosphoglycerate kinase 2 (PGK2),^59^ and phosphoglycerate mutase 1 and 2 (PGMA1/2).^60,61^ These enzymes exhibit pronounced hyperphosphorylation during capacitation, consistent with a switch-like mechanism that reactivates glycolysis. Notably, increased ALDOA activity was recently shown to drive the capacitation-induced rise in glycolytic flux,^62^ aligning with our phosphoproteomic findings. This reversible phospho-silencing likely limits ATP production and restricts premature motility, preserving energy stores until capacitation signals in the female tract relieve the metabolic brakes imposed during storage.

Coupled with the potential mechanisms of metabolic suppression described above, we also observed phosphoproteomic signatures related to the activation of protective mechanisms, which would also be compatible with promoting sperm longevity. For instance, we observed phosphorylation changes mapping to the NRF2-mediated oxidative stress response pathway and HIF1α signaling, both master regulators of antioxidant defenses.^63–65^ This suggests an adaptative mechanism for mitigating the constant threat of damage from reactive oxygen species produced by sperm mitochondria, thereby preserving the integrity of sperm DNA and membrane lipids essential for fertilization.^66,67^ Furthermore, our data highlights a potential role for protein homeostasis (proteostasis) pathways in maintaining sperm integrity during storage, with phosphoproteomic signatures consistent with activation of protein ubiquitination and the unfolded protein response occurring coincident with epididymal transit. ^68^ These quality control mechanisms presumably contribute to ensuring the retention of structurally and functionally sound proteins, while eliminating damage proteins that could otherwise impact cell survival and fertility. Consistent with this notion, we recorded increased phosphorylation of key elements of the ubiquitin-proteasome system, which is not only central to protein quality control networks, but also selectively targets proteins within the acrosome membrane for degradation, enabling controlled acrosome exocytosis during capacitation and fertilization.^69,70^ One such example is the evolutionarily conserved^31^ E1 ubiquitin-activating enzyme UBA1, which remains phosphorylated throughout sperm maturation. Through these coordinated phosphorylation events of metabolic, protective, and quality-control pathways, epididymal maturation successfully balances the dual demands of sperm storage and fertilization competence, priming sperm cells for the metabolic and functional demands of capacitation and fertilization, while maintaining them in a stable, quiescent state.

After leaving the epididymis and entering the female reproductive tract, spermatozoa undergo their final maturation phase known as capacitation, during which phosphorylation is finely tuned to activate essential fertilization functions.^3^ Our data revealed that while the overall number of phosphoproteomic changes accompanying capacitation is relatively modest compared to upstream epididymal transit, the functional impact of such changes is likely profound in terms of serving as a high-fidelity molecular switch to enable spermatozoa to realize their fertilization potential. Moreover, despite tyrosine phosphorylation traditionally being viewed as the primary driver of capacitation-related signaling events, we detected only a subtle increase in the overall proportion of phosphotyrosine residues identified in capacitated spermatozoa (i.e. up from 1.3% to 1.8% of phosphosites). Indeed, our analysis of significantly altered phosphosites during capacitation revealed that serine, threonine, and tyrosine residues were all proportionally targeted; with significant increases recorded against 10.2%, 8.2% and 9.4% of serine, threonine and tyrosine residues, respectively. This is exemplified by significant changes in phospho-serine and -threonine sites in key capacitation regulators such as cation channel sperm-associated proteins 2, 3, and 4 (CATSPER2/3/4),^71^ as well as the acrosome-associated protein ATP8B3.^72^ Such data demonstrates that capacitation is governed by a complex and coordinated symphony of kinase and phosphatase activities targeting all three amino acid types; a subtlety that has hitherto been overlooked using traditional antibody-based methods that lack the resolution of unbiased mass spectrometry approaches.

Occurring in concert with capacitation-associated phosphorylation events documented above, protein dephosphorylation events have also been implicated as playing critical roles in terms of reversing the metabolic “brakes” applied during epididymal maturation, and thereby allowing sperm to transition from a stable, storage-compatible state to an active, motile form capable of reaching and engaging with the oocyte.^73,74^ This coordinated phosphorylation and dephosphorylation of key proteins enable the sperm to respond effectively to environmental signals, preparing them for the demands of fertilization. For instance, we observed the simultaneous hyperphosphorylation of proteins in the RHOA signaling pathway, which is essential for effective acrosome reaction,^19,75^ and the dephosphorylation of proteins involved in inhibition of cytoskeleton organization, potentially releasing the cells from the inhibitory mechanisms needed to mitigate the risk of premature acrosomal exocytosis. This is further exemplified by the observation that proteins involved in binding of ova were hyperphosphorylated, while those linked to penetration of the zona pellucida were dephosphorylated, hinting at a sequential, exquisitely timed cascade of events that must occur for successful fertilization. The involvement of serine, threonine, and tyrosine amino acid targets in these changes suggests a more intricate network of phosphorylation controls than previously recognized. These findings demonstrate the critical importance of advanced phosphoproteomic approaches for understanding the biochemical complexities of capacitation and broaden our perspective on the regulatory mechanisms that prepare sperm for successful fertilization.

Beyond identification of site-specific phosphorylation events, this study provides a functional blueprint of the sperm kinome responsible for regulating these pathways, identifying a network of 343 kinases that putatively contribute to orchestrating the acquisition of fertilizing ability. This catalog represents an invaluable resource for the field, providing a comprehensive list of potentially druggable targets for manipulating fertility. Notably, several well characterized kinases involved in sperm capacitation were found amongst this resource, such as proto-oncogene tyrosine-protein kinase Src (SRC),^76,77^ tyrosine-protein kinase Fer (FER),^78^ and focal adhesion kinase 1 (FAK),^79^ highlighting the potential of this resource to uncover additional regulators of sperm function. Importantly, we moved beyond a descriptive inventory to provide direct functional validation of this resource. By focusing on kinases that become phosphorylated during capacitation, we identified serine/threonine-protein kinase 33 (STK33), homeodomain-interacting protein kinase 4 (HIPK4) and serine/threonine-protein kinase PAK 1 (PAK1) as novel regulators of sperm function. To test their roles, we used specific pharmacological inhibitors (ML281, Foretinib, and NVS-PAK1-1) at concentrations carefully optimized to avoid sperm toxicity and preserve viability. This strategy confirmed that the inhibition of each of these kinases significantly reduced the ability of sperm to undergo an ionophore-induced acrosome reaction. Strikingly, combined inhibition of all three kinase targets produced an additive effect, reducing acrosome exocytosis by 83%, effectively returning it to the basal levels observed in non-capacitated spermatozoa. Notably, since STK33 inhibition has previously been shown to be ineffective in reducing sperm ^29^, these data highlight the potential for different aspects of sperm function to be regulated by unique biochemical pathways..

To build on these observations, we examined the consequences of knocking out genes encoding proteins that underwent significant phosphorylation changes in our dataset through the European Mouse Mutant Archive (EMMA). ^28,31^ This analysis provided compelling *in vivo* validation of the functional importance of a subset of sperm proteins targeted for maturation related phosphorylation. For example, knockout of *Ccdc183a* resulted in a ∼27% reduction in sperm motility, while deletion of *Gsk3a* or *Optn* led to severe impairments in progressive motility, with reductions exceeding 34%. The effects on fertilization capacity were even more striking with sperm from *Aco2* knockout mice being completely incapable of fertilizing an egg, and those from *Cmpk1* knockouts exhibiting an 80.6% reduction in their ability to form two-cell embryos. The strong correlation between our identified phosphoproteins and these pronounced reproductive phenotypes offers powerful physiological support for the relevance of our findings and their future use by the field.

In establishing direct, causal links between the phosphorylation of specific kinases/proteins and the execution of critical fertilization events, our study provides a powerful proof-of-concept demonstration that (phospho)proteomic datasets can be harnessed to identify previously uncharacterized regulators of sperm competence. To facilitate seamless access to this resource, we developed and launched an interactive web application, ShinySpermPhopsho (https://reproproteomics.shinyapps.io/ShinySpermPhospho/) allowing researchers to explore the dataset and generate testable hypotheses for further investigation. By identifying these kinases and mapping their specific roles in sperm development, our findings contribute to a more comprehensive understanding of the regulatory networks driving sperm maturation. The phosphoproteomic insights described here illustrate the importance of kinase signaling in a spermatozoon’s journey to a fertilization competent cell, thereby providing new perspectives on potential therapeutic targets for male fertility treatments, as well as non-hormonal male contraceptives. Alternatively, the identification of specific phosphoproteins associated with sperm maturation offers up a range of valuable diagnostic biomarkers with the potential to inform the assessment of a male’s fertility status.

However, we also readily acknowledge the limitations in this study. In particular, the knockout mouse models used to validate the importance of a subset of phospho-targets were systemic rather than sperm-specific, meaning that we cannot fully exclude the potential contributions of developmental defects or hormonal imbalances to the observed phenotypes. Additionally, these models eliminate the entire protein rather than specifically modifying its phosphorylation state. Nonetheless, the fact that disruption of so many proteins identified in our phosphoproteomic screen leads to clear male subfertility phenotypes strongly supports the conclusion that phosphorylation-dependent regulation of these proteins is critical for normal reproductive function.

Overall, this study challenges traditional views on the role of phosphorylation in sperm biology, broadening the importance of phosphorylation beyond the regulation of capacitation and illustrating its crucial involvement in upstream post-testicular sperm maturation occurring within the context of epididymal maturation. The complex phosphoproteomic landscape uncovered here not only enhances our fundamental understanding of sperm function but also provides a foundation for future studies on male infertility and novel fertility treatments, ultimately improving reproductive health outcomes.

## ACKNOWLEDGEMENTS

We thank Dr Ben Crossett and Jens Altvater from the Mass Spectrometry Core Facility at The University of Sydney, and the Academic and Research Computing Support team, The University of Newcastle who provided High Performance Computing Infrastructure to support the bioinformatics analyses. We thank Steffie Dunst and Bernhard Rey for technical support with the sperm and IVF culture experiments. We thank the technicians and animal caretakers of the German Mouse Clinic. This research was supported by a National Health and Medical Research Council of Australia (NHMRC) Emerging Leadership Fellowship (APP2034392) awarded to D.A.S.B., Project Grant awarded to B.N. and M.D.D. (APP1147932). B.N., and M.D.D. are recipients of NHMRC Research Fellowships.

## AUTHOR CONTRIBUTIONS

Conceptualization, D.A.S.B. and B.N.; Methodology, D.A.S.B., S.J.H. and B.N.; Investigation, D.A.S.B., A.L.A., R.T., N.D.B., E.G.B., V.G.D., H.F., S.M., M.H.A., and S.J.H.; Formal Analysis D.A.S.B. and B.N.; Validation D.A.S.B., A.L.A., R.T., V.G.D., H.F., S.M., M.H.A. and B.N.,; Software, D.A.S.B.; Visualization, D.A.S.B., N.D.B., B.N.; Writing – Original Draft D.A.S.B. and B.N.; Writing – Review & Editing D.A.S.B., A.L.A, R.T., N.B., E.G.B., V.G.D., H.F., S.M., M.H.A., M.D.D., S.J.H. and B.N.; Funding Acquisition, D.A.S.B, M.D.D., and B.N.; Resources, S.J.H. and B.N.; Supervision, B.N.

## DECLARATION OF INTERESTS

The authors declare no competing interests.

## METHODS

### Data and Code Availability

- The mass spectrometry proteomics data have been deposited to the ProteomeXchange Consortium (http://proteomecentral.proteomexchange.org) via the PRIDE partner repository ^81^ with the dataset identifier PXD041286 and 10.6019/PXD041286. PRIDE Reviewer account details: Username; Password: YJq9oPpm
- All original code has been deposited at GitHub and is publicly available at https://github.com/DavidSBEire/ShinySpermPhopsho as of the date of publication.
- Any additional information required to reanalyze the data reported in this paper is available from the corresponding authors upon request.

### Ethics approval

All experimental procedures involving mice were conducted with the approval of the University of Newcastle (UoN) Animal Care and Ethics Committee (ACEC; approval number A-2018-826). Male outbred Swiss mice were obtained from a breeding colony held at the UoN central animal facility and maintained according to the recommendations prescribed by the ACEC. Mice were housed under a controlled lighting regimen (12L:12D) at 21°C – 22°C and supplied with food and water *ad libitum*.

### Isolation of epididymal spermatozoa

Immediately after adult male mice (8 week old; n = 6 / biological replicate) were euthanized, their vasculature was perfused with pre-warmed Tris-buffered saline (TBS) to minimize the possibility of blood contamination. The epididymides were then removed, separated from fat and overlying connective tissue, and carefully dissected to isolate two anatomical segments of interest; the caput (proximal segment) and cauda (distal segment). The caput spermatozoa were recovered by placing the tissue in a 500 μl droplet of modified Biggers, Whitten, and Whittingham media (BWW; ^19,82^) composed of 91.5 mM NaCl, 4.6 mM KCl, 1.7 mM CaCl_2_·2H_2_O, 1.2 mM KH_2_PO_4_, 1.2 mM MgSO_4_·7H_2_O, 25 mM NaHCO_3_, 5.6 mM D-glucose, 0.27 mM sodium pyruvate, 44 mM sodium lactate, 5 U/ml penicillin, 5 μg/ml streptomycin, 20 mM Hepes buffer, and 3 mg/ml bovine serum albumin [BSA]) (pH 7.4; osmolarity 300 mOsm/kg). After multiple incisions were made with a razor blade, the spermatozoa were gently washed into the medium with mild agitation. All sperm preparations were passed through a 70 µm filter, then subjected to centrifugation (400 × g for 15 min) on a 28% Percoll/BWW density gradient. The pellet, consisting of an enriched population of caput spermatozoa (Figure S1A), was resuspended in BWW/PVA (BWW as above, with the exception that 1 mg/mL polyvinyl alcohol (PVA) was substituted in place of 3 mg/mL BSA) and re-centrifuged (400 × g for 2 min) to remove excess Percoll. Notably, BWW supplementation with BSA supports significantly higher rates of *in vitro* sperm capacitation (both in terms of the time taken to capacitate and the percentage of sperm that achieve capacitation) ^11^, yet the replacement with PVA in the BWW wash buffer is designed to eliminate residual BSA prior to cell lysis and downstream applications involving protein analysis ^83^. The purity and vitality of each sperm preparation was confirmed by microscopy (purity > 95% spermatozoa and > 70% motility) consistent with previous studies^19,84–86^.

Cauda epididymal spermatozoa were collected from the lumen via retrograde perfusion with water-saturated paraffin oil as previously described ^83^. To prepare non-capacitated (NC Cauda) and capacitated (CAP Cauda) populations of cauda spermatozoa, the following medias/conditions were used. Spermatozoa were held in a non-capacitated state in NC-BWW (BWW prepared as described above with the exception that 25mM NaCl was substituted in place of the 25mM NaHCO_3_), and prepared immediately after their collection. By contrast, capacitated spermatozoa were driven to capacitate in BWW supplemented with 1 mM pentoxifylline (ptx) and 1 mM dibutyryl cyclic adenosine monophosphate (dbcAMP) and incubated at a concentration of ∼10 million/mL at 37°C 5% CO_2_ for 45 min (with tube lids loosened permissive of gas exchange) ^19,82,83^. Spermatozoa were washed in BWW/PVA media after capacitation to remove any residual BSA before snap freezing and prepared for proteomic analysis as described below.

### SDS-PAGE and Immunoblotting

Sperm proteins were extracted from samples independent of those used for mass spectrometry using a modified SDS-PAGE sample buffer (2% w/v SDS, 10% w/v sucrose in 0.1875 M Tris, pH 6.8) supplemented with protease inhibitor cocktail (Roche, Basel, Switzerland). Samples were boiled at 100°C for 5 min after which insoluble matter was pelleted by centrifugation at 20,000 × g for 10 min and the quantity of soluble protein remaining in the supernatant was estimated using a DC Protein Assay in accordance with the manufacturer’s instructions (Bio-Rad Laboratories, Hercules, CA). Similarly, proteins were also extracted using different methods such as RIPA and SDC buffers. For RIPA buffer (150 mM NaCl, 1% v/v Triton X-100, 0.5% w/v sodium deoxycholate, 0.1% w/v SDS, 50 mM Tris pH 8) the protein was shaken at 4°C for 20 min, and for SDC (4% w/v SDC and 100 mM Tris-HCl pH8.5), protein was sonicated 4 ×10s. Both cell lysate suspensions were then centrifuged, and protein quantified as above. Extracted proteins were boiled for 5 min in SDS-PAGE sample buffer (2% v/v mercaptoethanol, 2% w/v SDS, and 10% w/v sucrose in 0.1875 M Tris, pH 6.8, with bromophenol blue) prior to being resolved by SDS-PAGE and transferred onto nitrocellulose membranes. Membranes were blocked with 3% w/v bovine serum albumin (BSA) in TBS and 0.1% v/v polyoxyethylenesorbitan monolaurate (Tween-20; TBS-T; pH 7.4) for 1 h before being probed with 1 µg/mL concentration of appropriate primary antibodies diluted in TBS-T containing 1% w/v BSA overnight at 4°C. Blots were washed three times in TBS-T followed by incubation with an appropriate horseradish peroxidase (HRP)-conjugated secondary antibodies diluted 1:2500 in 1% w/v BSA/TBS-T for 1 h at room temperature. Following three washes in TBS-T, labeled proteins were detected using an enhanced chemiluminescence kit (GE Healthcare, Chicago, IL) and visualized on ChemiDoc MP (Bio-Rad, Hercules, California, USA) ^19,87,88^. For the validation of each candidate protein, three immunoblots were performed using three independent biological samples, with each blot being subjected to stripping and re-probing with the loading control α-GAPDH antibody to in preparation for densitometric analyses of relative band labeling intensity.

### Immunofluorescence

Spermatozoa were fixed in 4% PFA, washed in 50 mM glycine/ phosphate-buffered saline (PBS), and settled onto poly-L-lysine-treated coverslips at 4°C overnight. All subsequent incubations were performed in a humidified chamber, and all antibody dilutions and washes were conducted in PBS. Fixed cells were permeabilized in 0.2% Triton X-100/PBS for 10 min and blocked in 3% (w/v) BSA in PBS for 1 h at 37°C. Slides were then sequentially labeled with primary antibodies (diluted between 1:50 to 1:250) overnight at 4°C. After incubation, the slides were washed three times, then incubated in appropriate 488 Alexa Fluor conjugated secondary antibodies (diluted 1:400) for 1 h at 37°C. Cells were then washed and mounted in antifade reagent (Mowiol 4-88). Labeled cells were viewed on an Axio Imager A2 microscope (Carl Zeiss MicroImaging, Inc., Jena, Germany) equipped with epifluorescent optics and images captured with Zeiss Axiocam 305 mono camera.

### Phosphoproteomic sample preparation of spermatozoa

All samples were prepared for phosphoproteomic analysis using methodology known as EasyPhos (EP) ^19,20,89^. In brief, 250 µL of chilled lysis buffer [4% (w/v) sodium deoxycholate (SDC); 100 mM Tris-HCl (pH 8.5)], was added to each sample and immediately heated (95°C, 5 min) to inactivate endogenous proteases and phosphatases. Samples were sonicated (4 × 20 s cycles, 75% output power), and an aliquot taken to determine protein concentration using a bicinchoninic acid assay (BCA). All samples were diluted to equal protein amounts (120 µg) in 270 µL of lysis buffer into a 2 mL 96 deepwell plate. Samples were reduced and alkylated [100 mM Tris(2-carboxyethyl)phosphine hydrochloride; 400 mM 2-chloroacetemide], using a Thermomixer (Eppendorf; Hamburg, Germany), samples were incubated for 5 min at 45°C (1,500 rpm). Enzymatic digestion was achieved using Lys-C and trypsin, at an enzyme-to-substrate ratio of 1:100 (w/w) and incubated overnight at 37°C with shaking (1,500 rpm). To the digested peptides, 400µL of isopropanol (ISO) was added and mixed thoroughly for 30 s on the Thermomixer (1,500 rpm). Next 100µL of EP Enrichment buffer [48% (v/v) TFA / 8 mM potassium dihydrogen phosphate (KH2PO4)] was added to each sample and mixed for 30 s thoroughly (1,500 rpm). The plate containing the peptides was spun at 2,000 x g for 15 min (RT) and supernatants carefully transfer to a new clean 2 mL 96 deepwell plate. To each sample, resuspend in EP loading buffer [6% (v/v) TFA / 80% (v/v) ACN; concentration of mg / μl^−1^] 5mg of TiO_2_ beads were added and incubated at 40°C with shaking (2,000 rpm). Beads were pelleted (2,000 x g, 1 min) and the supernatant removed. The beads were washed five times with 1 mL EP wash buffer [5% (v/v) TFA / 60% (v/v) ISO], each mixed briefly (1,500 rpm) and supernatant removed after spinning (2,000 x g, 1 min). Beads were resuspended in 75 µL of EP transfer buffer [0.1% (v/v) TFA / 60% (vol/vol) ISO] and transferred to a C8 StageTip ^90^. An additional 75 µL EP transfer buffer was added to each well to ensure all beads were retained and transferred to their C8 StageTip. To an in-house 3D printed 96-well StageTip centrifuge device ^19,20^, each StageTip was secured and spun through to dryness (1,500 x g, 8 min). Next, 2 x 60µL of EP elution buffer [200 μl of 25% ammonia solution / 800 μl of 40% (v/v) ACN.] was added to each StageTip and spun in PCR tubes (1,500 x g, 3 min). Immediately, the PCR tubes were lyophilized using a centrifuge concentrator (Eppendorf; Hamburg, Germany) at 45°C for 30 minutes, ensuring the samples are not completely dry (∼20 µL). To the partially dried samples, 100µL of styrenedivinylbenzene-reverse phase sulfonated (SDB-RPS) loading buffer [1% (v/v) TFA in ISO] was added and each sample transferred to a SDB-RPS StageTip ^90^ for desalting ^19^. StageTips were centrifuged at 1,500 × g for 3 min and subsequently washed with 100µL 99% ethylacetate/1% TFA (1,500 × g, 3 min). Finally, StageTips were washed with 100 µL each of 99% ISO/1% TFA and 0.2% TFA/5% acetonitrile, eluted with 60 µL of 5% NH_4_OH/80% acetonitrile, dried by vacuum concentration and re-suspended in MS loading buffer (2% acetonitrile/0.3% TFA). Purified sperm phosphopeptides were subjected to analysis by high resolution nano liquid chromatography tandem mass spectrometry (nLC-MS/MS).

### nLC-MS/MS analysis

Reverse phase nLC-MS/MS was performed using an Orbitrap HF-X MS coupled to a Dionex Ultimate 3000RSLC nanoflow high-performance liquid chromatography system (Thermo Fisher Scientific). Peptide separation was then achieved using an in-house packed column, SGE MyCapLC Kit (Kinesis) 300 mm x 150 mm, employing an 87 min stepped gradient of acetonitrile (0.350 µL/min; 3 - 20%, 52 min; 20 - 40%, 20 min; 40% - 98%, 15 min). Full MS/data dependent acquisition MS/MS mode was utilized on Xcalibur (Thermo Fisher Scientific; version 4.2.47) with the Orbitrap mass analyzer set at a resolution of 60,000, to acquire full MS with a range of 350 – 1400 m/z, incorporating an automatic gain control target of 3 x 10^6^ and maximum fill times of 120 ms. The 10 most intense multiply charged precursors were selected for higher-energy collision dissociation fragmentation with a normalized collisional energy of 27. MS/MS fragments were measured at an Orbitrap resolution of 15,000 using an automatic gain control target of 1 x 10^5^ and maximum fill times of 50 ms ^20^.

### Kinase inhibition studies

Initial cytotoxicity testing was undertaken whereby cauda epididymal spermatozoa were dispersed into NC-BWW and separated into equal populations prior to the addition of inhibitors for either STK33 (ML281), HIPK4 (Foretinib) and PAK1 (NVS-PAK1-1) at increase concentrations or an equal volume of BWW medium (control). The spermatozoa were pre-incubated with or without inhibitor for 0, 60 min at 37°C, before sperm viability was assessed (Figure S4). Additionally, motility and progressive motility were assessed using the computer-assisted sperm analysis (CASA) system.^91^ Following this, cauda epididymal spermatozoa were dispersed into NC-BWW and separated into two equal populations prior to the addition of the inhibitor (maximum concentration) to the treatment group and an equal volume of medium to the control group. The spermatozoa were pre-incubated with or without inhibitor for 1 h at 37°C, before being driven to capacitate (as described above) for a further 45 min at 37°C in an atmosphere of 5% CO_2_. At the end of each incubation, the sperm populations were assessed for motility and were then either fixed in 4% paraformaldehyde (for assessment of spontaneous acrosome reaction rates) or induced to undergo an acrosome reaction with the calcium ionophore, A23187 (Merck) (induced). For the latter treatment, A23187 was added at a final concentration of 2.5 µM, alongside a DMSO vehicle control, and incubated 37°C for 30 min. After induction of acrosomal exocytosis, spermatozoa were incubated in hypoosmotic swelling medium for 1 h at 37°C and then fixed in 4% paraformaldehyde. Spermatozoa were then washed in 50 mM glycine/PBS and air dried onto 12-well slides in preparation for labeling with the acrosome marker, PNA-FITC (lectin from *Arachis hypogea* [peanut]) (Merck). Spermatozoa were first permeabilized with ice cold methanol for 10 min followed by incubation in PNA-FITC (5 µg/mL) for 20 min at 37°C, and 3 final washes in PBS. Spermatozoa were examined using a Zeiss Axio Imager 2 and the percentage of acrosome reacted cells (i.e. those without florescent labeling of their acrosome) was calculated ^19,92^.

### International Mouse Phenotyping Consortium mouse models, histology, and data collection

The International Mouse Phenotyping Consortium (IMPC) database ^93,94^ was mined for genetic knockout mice overlapping with those identified phosphorylated proteins altered through epididymal transit or capacitation and with special access granted, we crossed referenced with the European Mouse Mutant Archive (EMMA) ^28^ to restrict to those gene KOs with available *in vitro* fertilization (IVF) and sperm data. The mouse models were generated using the IMPC targeting strategy with CRISPR/Cas technology at Helmholtz Munich (https://www.mousephenotype.org/understand/the-data/allele-design/). After genotyping, heterozygous × heterozygous matings were set up to generate sufficient mutant mice with littermate ^+/+^ controls for phenotyping analysis at the German Mouse Clinic as described.^95^ We obtained data pertinent to *Aco2* (Aconitate hydratase, mitochondrial), Ahctf1 (Protein ELYS), *Atp8b3* (Phospholipid-transporting ATPase IK), *Cap2* (Adenylyl cyclase-associated protein 2), *Ccdc183* (Coiled-coil domain-containing protein 183), *Cenpv* (Centromere protein V), *Cfap47* (Cilia and flagella-associated protein 47), *Cmpk*1 (UMP-CMP kinase), *Dhx9* (Isoform 2 of ATP-dependent RNA helicase A), *Gdi2* (Rab GDP dissociation inhibitor beta), *Gsk3a* (Glycogen synthase kinase-3 alpha), *Gstm5* (Glutathione S-transferase Mu 5), *Lpp* (Isoform 4 of Lipoma-preferred partner homolog), *Nt5c2* (Cytosolic purine 5’-nucleotidase), *Nup35* (Nucleoporin NUP35), *Optn* (Optineurin), *Pbrm1* (Protein polybromo-1), *Ppil6* (Probable inactive peptidyl-prolyl cis-trans isomerase-like 6), and *Rps26* (40S ribosomal protein S26), with a wildtype reference established by EMMA.^31^ For further information on protocols used by EMMA for sperm collection, analysis and IVF, please see their publicly available resources and videos (https://www.infrafrontier.eu/emma/cryopreservation-protocols/).

### Immunoblot densitometry

Image Lab software (Bio-Rad) was used to perform densitometric analysis of all immunoblots. Briefly, each lane was labeled and the correct sized band of the protein of interest was marked. The ‘lane profile’ tool was used to define the correct lane and band size widths and to confirm uniformity. An ‘analysis table’ was then generated to contain the densitometric volume for each band in each lane (adjusted by subtraction of background). This process was replicated with the corresponding α-GAPDH immunoblot image to enable the volume of the band of interest to normalized against that of the comparable α-GAPDH loading control band volume for each lane. The fold change of band volume from cauda sperm samples was calculated relative to that of the caput sperm samples, thereby enabling determination of changes in the relative abundance of each protein of interest. Three such analyses were conducted on independent biological samples for each protein of interest.

### Proteomic data processing and analysis

Consistent with previous studies ^19,87,89,96–99^, database searching of raw files were performed separately using Proteome Discoverer 2.5 (Thermo Fisher Scientific). SEQUEST HT was used to search against the UniProt *Mus musculus* database (25,424 sequences, downloaded 21^st^ June 2022). Database searching parameters included up to two missed cleavages, a precursor mass tolerance set to 10 ppm and fragment mass tolerance of 0.02 Da. Trypsin was designated as the digestion enzyme. Cysteine carbamidomethylation was set as a fixed modification while phosphorylation (S, T, Y) was designated as a dynamic modification. Interrogation of the corresponding reversed database was also performed to evaluate the false discovery rate (FDR) of peptide identification using Percolator on the basis of q-values, which were estimated from the target-decoy search approach. To filter out target peptide spectrum matches over the decoy-peptide spectrum matches, a fixed FDR of 1% was set at the peptide level. Label-free quantification was performed using Proteome Discoverer nodes “Minora Feature Detector”, “Feature Mapper”, and “Precursor Ions Quantifier” nodes as described previously ^87^. Fold change and significance (*t*-test) comparative testing between sample groups was carried out within the Proteome Discoverer 2.5 suite. The peptide list was exported from Proteome Discoverer 2.5 as an Excel file and further refined to include only those with a quantitative value in all replicates of at least one of the sample groups. The refined peptide list was loaded into Perseus, version 1.6.10.43 ^100^, for the generation of scatter plots, principal component analysis, heatmaps, and ANOVA statistical testing. Basic data handling, if not otherwise stated, was conducted using Microsoft Excel 365 (Version 2211, Microsoft Corporation, Redmond, WA) and GraphPad Prism version 9.4.1 for Windows (GraphPad Software; San Diego, CA).

### Ingenuity pathway analysis

For each of the sample groups, where a phosphopeptide was expressed in all replicates, the corresponding UniProt accession numbers and transformed ratios were analyzed using Ingenuity Pathway Analysis software (IPA®, Qiagen) as previously described ^19,87,99,101,102^. For caput (2,293/2,299; 99.7%), non-capacitated cauda (1,373/1,377; 99.7%) and capacitated cauda (1,339/1,343; 99.7%) sperm phosphoproteomes were analyzed on the basis of predicted subcellular location and classification (other excluded). Importantly, the IPA phosphorylation specific knowledge base was utilized, ensuring the biological predictions are based upon patterns of phosphorylation changes and not protein expression. For ANOVA significant group analyses, a list for each group containing UniProt accession numbers and transformed ratios were submitted to IPA. Canonical pathway, upstream regulators, and disease and function analyses were assessed using; *p*-value, an enrichment measurement of the overlapping proteins from the dataset in a particular pathway, function or regulator ^103^ and; *Z*-score, a prediction scoring system of activation or inhibition based upon statistically significant patterns in the dataset and prior biological knowledge manually curated in the Ingenuity Knowledge Base. To elucidate the most significant changes in our analyses, we applied a stringency criteria of -log10 *p*-value of > 1.3 for each timepoint, and a Z-score of (inhibition) −2 ≤ Z ≥ 2 (activated) in at least one group ^19,87,99,104^. For disease and function assessment we restricted the analysis to ‘molecular and cellular functions’ and ‘physiological system development and function’. For assessment of the temporal groups, the top ten most significantly enriched overlapping canonical pathways and molecular and cellular functions were reported.

### Kinase identification

In addition to the direct identification of kinases within the phosphoproteomic dataset and the upstream regulator feature of IPA, the dataset was subjected to a kinase-substrate analysis utilizing the PhosphoSitePlus database (accessed 5^th^ May 2021, http://www.phosphosite.org/), an open-source curated resource for investigating the importance of experimentally observed post-translational modifications in the regulation of biological processes ^27^, as previously described.^26^ KinMap ^80^ was utilized in the visual representation of the findings.

### Shiny Application development

In accordance with Shiny blueprint outlined by ShinySperm,^47^ ShinySpermKingdom,^105^ and ShinySpermPlacenta,^23^ a Shiny Application was deployed to support the accessibility and interpretability of these datasets within, allowing for effective data-driven insights by the field. The full coding script supporting ShinySpermPhospho (https://reproproteomics.shinyapps.io/ShinySpermPhospho/), can be downloaded from GitHub – https://github.com/DavidSBEire/ShinySpermPhospho. In brief, the ShinySpermPhospho application was built using the shiny package (version 1.9.1) on RStudio (version 2024.04.1+748), with base *R* (version 4.3.3, 2024-02-29). Supporting the functionality and aesthetics of this application are several packages, including: DT, eulerr, ggplot2, openxlsx, plotly, readxl, reshape2, RColorBrewer, and shinydashboard.

### Statistical Analysis

Phosphoproteomic analyses were performed using sperm cell populations collected from two anatomical segments of interest; the caput (proximal region) and cauda (distal region) with a portion of caudal spermatozoa induced to undergo capacitation *in vitro* as described above (n = 3 biological replicates collected from 6 individual mice/biological replicate). Differentially accumulated sperm proteins were defined as those with a fold-change ± 1.5 and *p-*value ≤ 0.05. All other data were assessed for normality using a Shapiro-Wilk normality test. Normally distributed data were analyzed by unpaired Student’s t-tests to detect differences between treatment groups. Data not normally distributed were analyzed by a Mann-Whitney test. Differences between groups were considered significant when *p* ≤ 0.05. The number of biological replicates used in each experiment are presented in figure captions. Graphical data were prepared using GraphPad Prism (version 10.0.0) and are presented as mean values ± SEM.

## SUPPLEMENTAL FIGURES

**Figure S1:**
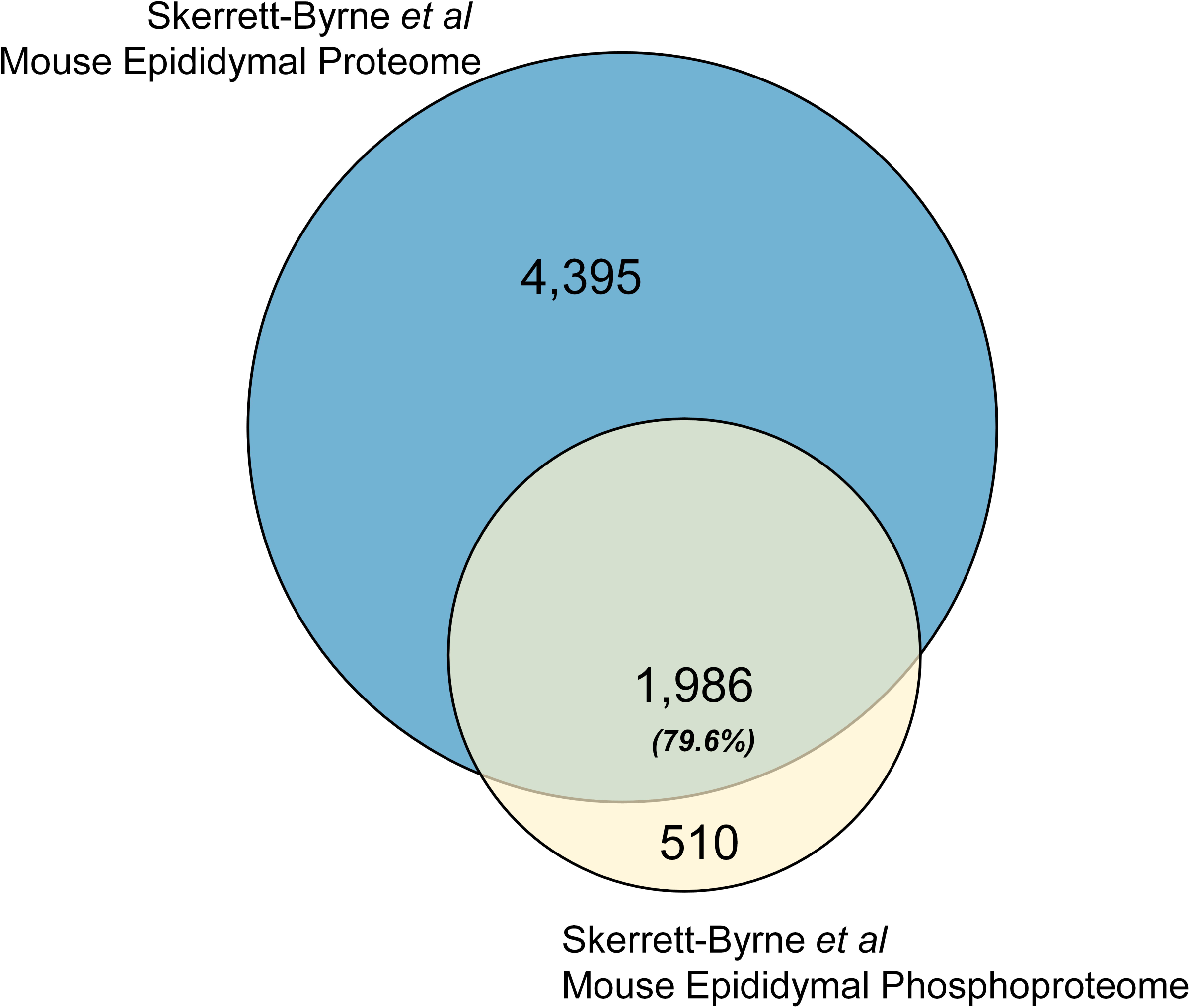
Comparison between epididymal sperm proteome and phosphoproteome. A direct comparison of the total proteins identified in ^19^ mouse epididymal sperm proteome and the phosphoproteome herein.

**Figure S2:**
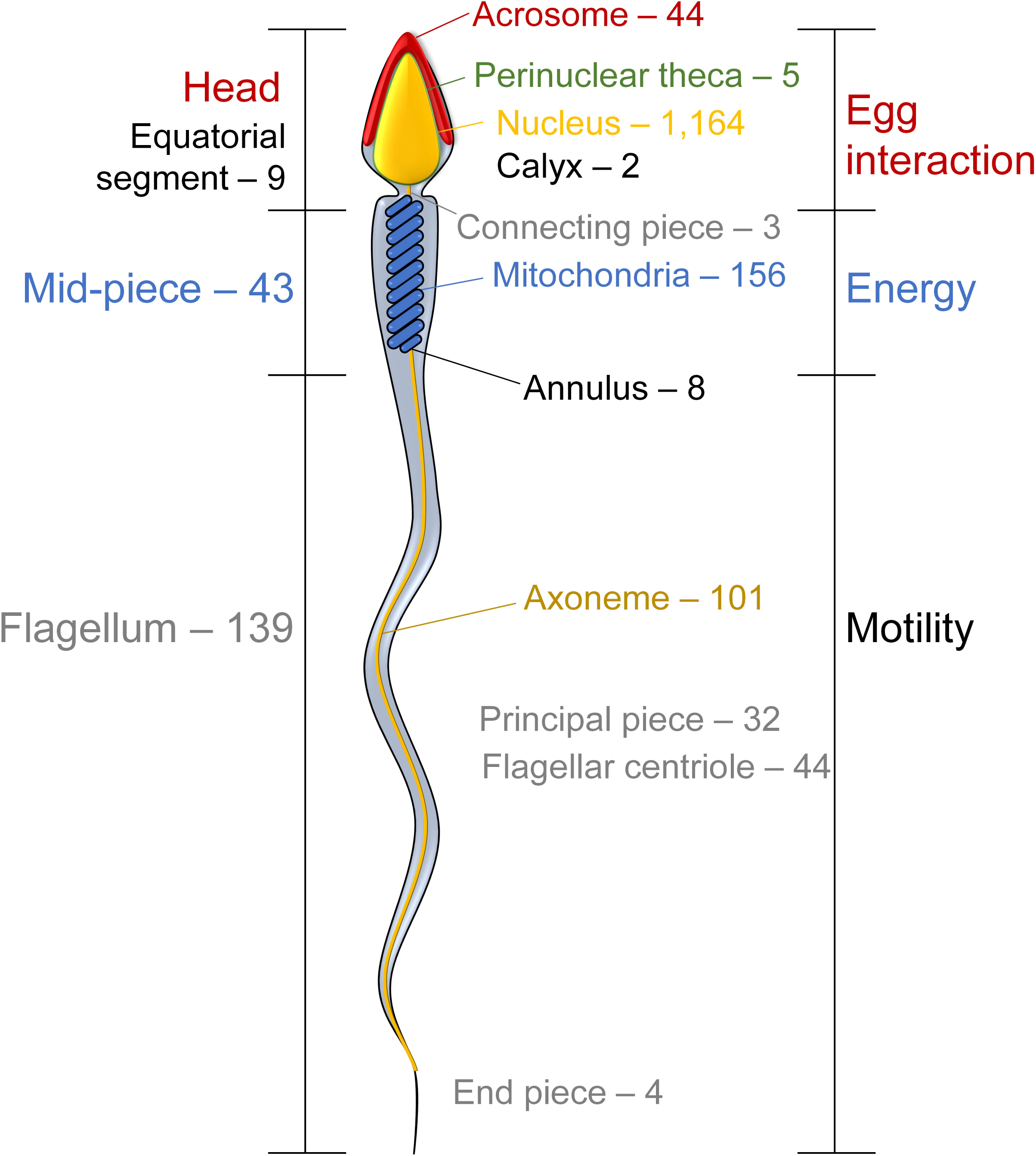
Sperm phosphoprotein cellular location. *In-silico* localization of identified phosphoproteins, mapping to key sperm regions; acrosome, annulus, axoneme, flagellum, mid-piece, mitochondria, nucleus and perinuclear theca, using UniProt and EMBL-EBI GO annotations. See Tables S1-4.

**Figure S3:**
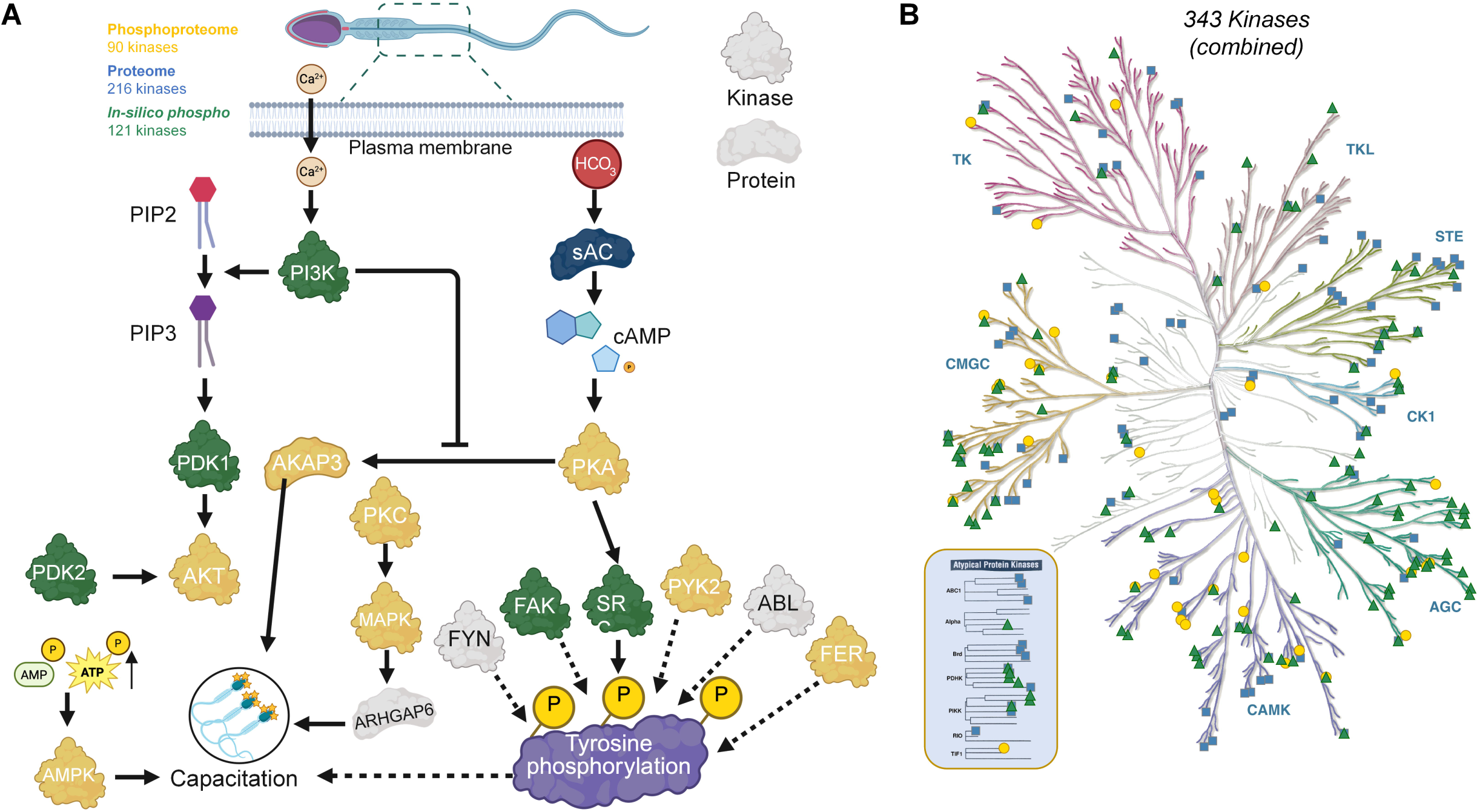
Summary of identified kinases and families. (A) All kinase identified in phosphoproteome dataset, epididymal sperm proteome and *in-silico* analyses, mapped onto the current knowledge of kinases and proteins regulating sperm capacitation and tyrosine phosphorylation. (B) Using KinMap,^80^ the combined 343 kinases (duplicates across datasets removed) are mapped on the Kinome map, into the eight typical kinase families groups and 13 atypical families; Protein Kinase A, G, and C families (AGC), Calcium/Calmodulin-Dependent Protein Kinase family (CAMK), Casein Kinase 1 family (CK1), Cyclin-Dependent Kinase, MAP Kinase, GSK3, and CDK-like family (CMGC), Sterile Kinase family (STE), Tyrosine Kinase family (TK), Tyrosine Kinase-Like family (TKL) and Other. Gold circles (phosphoproteome), blue squares (proteome),^19^ and green triangles (*in-silico* analyses). See Table S6.

**Figure S4:**
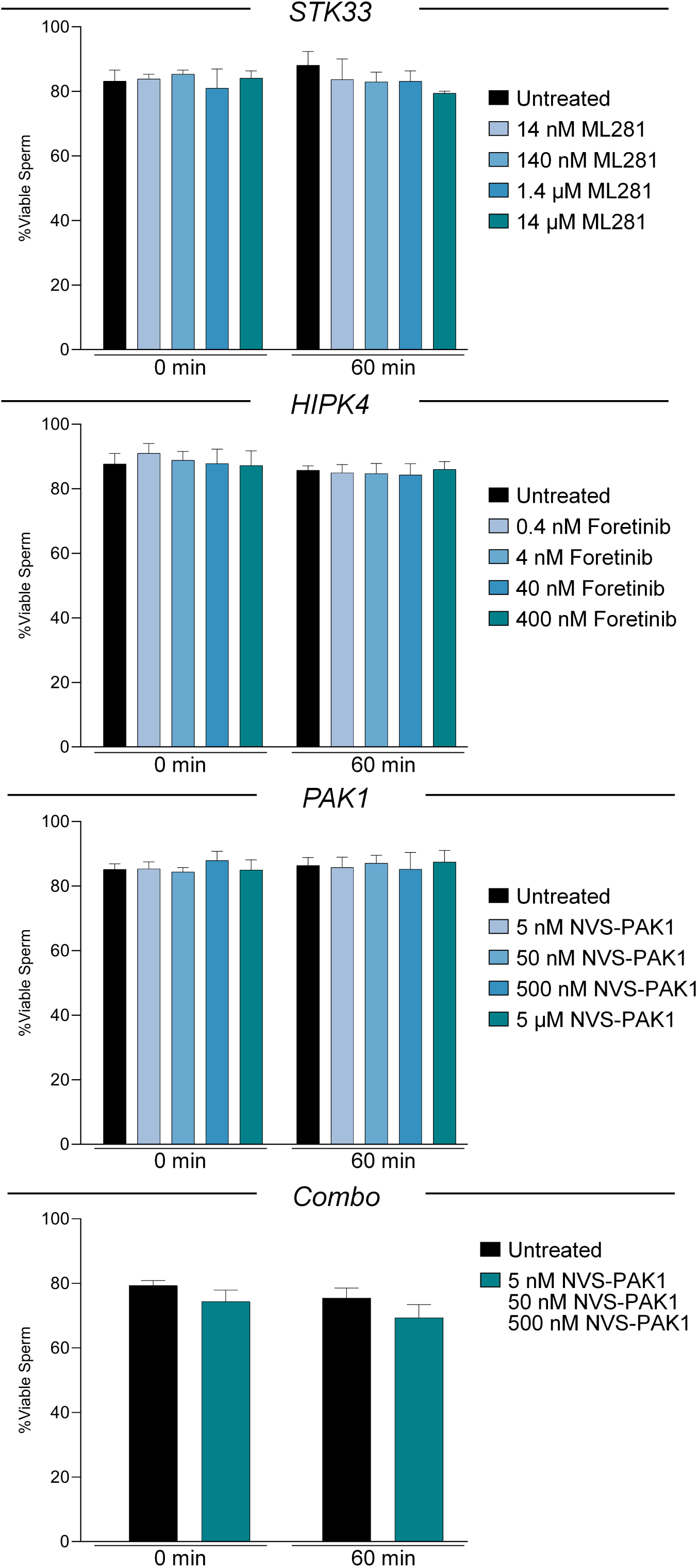
Cytotoxicity assessment on sperm viability. Mature cauda spermatozoa were assessed for potential cytotoxicity of each inhibitor alone and in combination; STK33 (ML281), HIPK4 (Foretinib) and PAK1 (NVS-PAK1-1). Alongside an untreated sample, four concentrations of each inhibitor were tested following a 1 hr treatment, whereby the number of viable sperm were counted.

**Figure S5:**
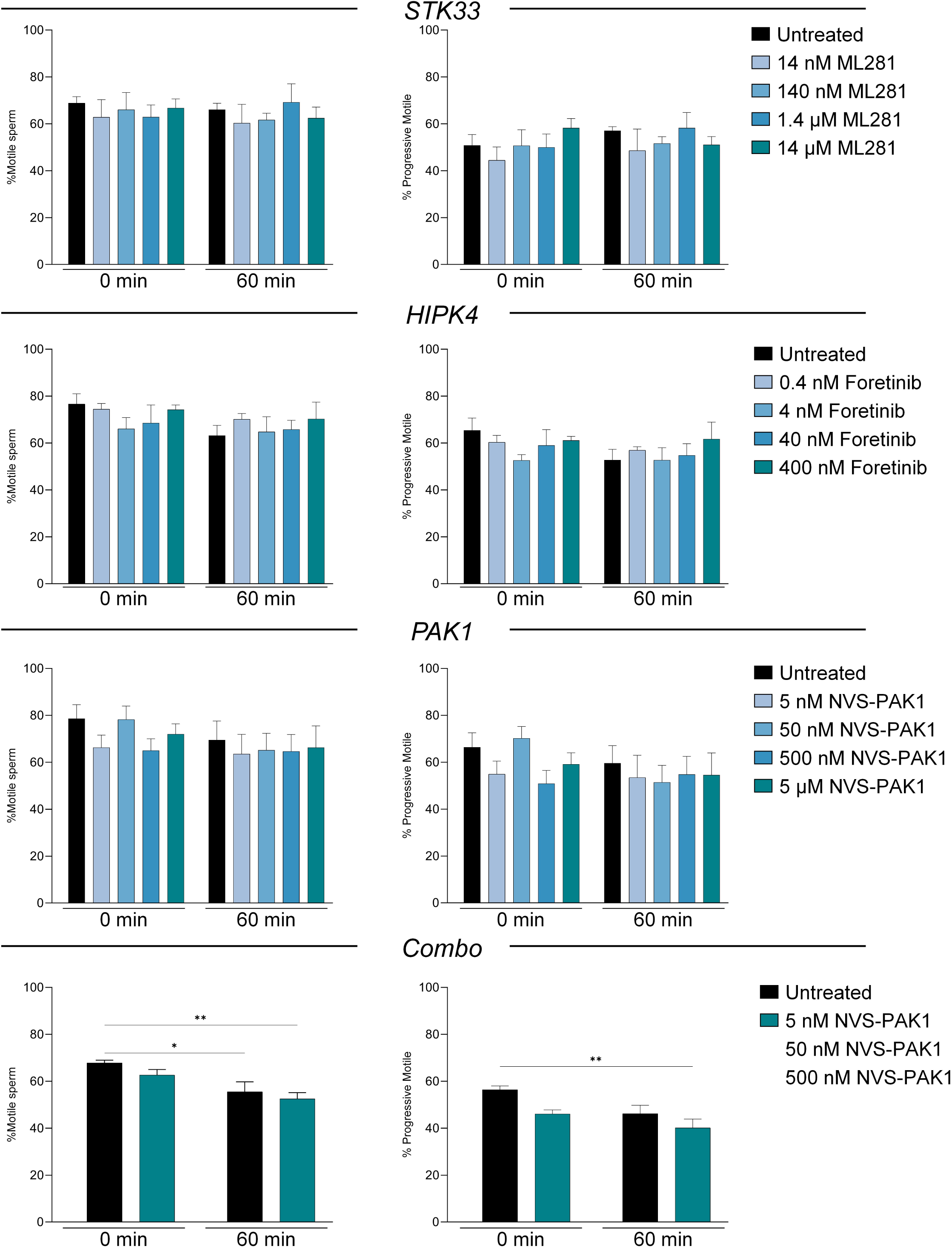
Cytotoxicity assessment on sperm motility and progressive motility. Four increasing concentrations of each inhibitor were assessed for their potential impact on sperm motility and progressive motility, in addition to an untreated control, using Computer Assisted Semen Analysis (CASA); STK33 (ML281), HIPK4 (Foretinib) and PAK1 (NVS-PAK1-1).

## SUPPLEMENTAL TABLES

**Table S1** Full phosphoproteomic characterization of the functionally immature caput epididymal spermatozoa. Data related to Figures 2 and 3.

**Table S2** Full phosphoproteomic characterization of the functionally mature caput epididymal spermatozoa (NC). Data related to Figures 2 and 3.

**Table S3** Full phosphoproteomic characterization of the functionally mature caput epididymal spermatozoa (CAP) which have undergone capacitation *in vitro* prior to cell lysis. Data related to Figures 2 and 3.

**Table S4** The ANOVA significant phosphopeptides with group numbers annotated. Data related to Figure 4.

**Table S5** Complete comparative phosphoproteomic analyses between all three epididymal spermatozoa (caput, NC, and CAP). Data related to Figures 2, 3, and 5.

**Table S6** All kinase identified in phosphoproteome data and *in-silico* analyses. Data related to Figure 6 and S3, and the Ingenuity pathway analysis and kinase mapping section in the STAR Methods.

**Table S7** Specific phosphopeptide data mapping to those proteins explored through knock-out mouse models

## Notes

https://reproproteomics.shinyapps.io/ShinySpermPhospho/

### Competing Interest Statement

The authors have declared no competing interest.

## REFERENCES

1. Nixon, B., Cafe, S.L., Eamens, A.L., De Iuliis, G.N., Bromfield, E.G., Martin, J.H., Skerrett-Byrne, D.A., and Dun, M.D. (2020). Molecular insights into the divergence and diversity of post-testicular maturation strategies. Mol Cell Endocrinol 517, 110955. 10.1016/j.mce.2020.110955.

2. Zhou, W., De Iuliis, G.N., Dun, M.D., and Nixon, B. (2018). Characteristics of the Epididymal Luminal Environment Responsible for Sperm Maturation and Storage. Front Endocrinol (Lausanne) 9, 59. 10.3389/fendo.2018.00059.

3. Aitken, R.J., and Nixon, B. (2013). Sperm capacitation: a distant landscape glimpsed but unexplored. Mol Hum Reprod 19, 785–793. 10.1093/molehr/gat067.

4. Aitken, R.J., Nixon, B., Lin, M., Koppers, A.J., Lee, Y.H., and Baker, M.A. (2007). Proteomic changes in mammalian spermatozoa during epididymal maturation. Asian J Androl 9, 554–564. 10.1111/j.1745-7262.2007.00280.x.

5. Baker, M.A., Nixon, B., Naumovski, N., and Aitken, R.J. (2012). Proteomic insights into the maturation and capacitation of mammalian spermatozoa. Syst Biol Reprod Med 58, 211–217. 10.3109/19396368.2011.639844.

6. Dacheux, J.L., Belleannee, C., Jones, R., Labas, V., Belghazi, M., Guyonnet, B., Druart, X., Gatti, J.L., and Dacheux, F. (2009). Mammalian epididymal proteome. Mol Cell Endocrinol 306, 45–50. 10.1016/j.mce.2009.03.007.

7. Gatti, J.L., Castella, S., Dacheux, F., Ecroyd, H., Metayer, S., Thimon, V., and Dacheux, J.L. (2004). Post-testicular sperm environment and fertility. Anim Reprod Sci 82-83, 321-339. 10.1016/j.anireprosci.2004.05.011.

8. Labas, V., Spina, L., Belleannee, C., Teixeira-Gomes, A.P., Gargaros, A., Dacheux, F., and Dacheux, J.L. (2015). Analysis of epididymal sperm maturation by MALDI profiling and top-down mass spectrometry. J Proteomics 113, 226–243. 10.1016/j.jprot.2014.09.031.

9. Skerrett-Byrne, D.A., and Nixon, B. (2022). Cell Reports.

10. Walsh, C.T., Garneau-Tsodikova, S., and Gatto, G.J., Jr. (2005). Protein posttranslational modifications: the chemistry of proteome diversifications. Angew Chem Int Ed Engl 44, 7342–7372. 10.1002/anie.200501023.

11. Asquith, K.L., Baleato, R.M., McLaughlin, E.A., Nixon, B., and Aitken, R.J. (2004). Tyrosine phosphorylation activates surface chaperones facilitating sperm-zona recognition. J Cell Sci 117, 3645–3657. 10.1242/jcs.01214.

12. Balbach, M., Gervasi, M.G., Hidalgo, D.M., Visconti, P.E., Levin, L.R., and Buck, J. (2020). Metabolic changes in mouse sperm during capacitationdagger. Biol Reprod 103, 791–801. 10.1093/biolre/ioaa114.

13. Nixon, B., Bielanowicz, A., Anderson, A.L., Walsh, A., Hall, T., McCloghry, A., and Aitken, R.J. (2010). Elucidation of the signaling pathways that underpin capacitation-associated surface phosphotyrosine expression in mouse spermatozoa. J Cell Physiol 224, 71–83. 10.1002/jcp.22090.

14. Porambo, J.R., Salicioni, A.M., Visconti, P.E., and Platt, M.D. (2012). Sperm phosphoproteomics: historical perspectives and current methodologies. Expert Rev Proteomics 9, 533–548. 10.1586/epr.12.41.

15. Stival, C., Puga Molina Ldel, C., Paudel, B., Buffone, M.G., Visconti, P.E., and Krapf, D. (2016). Sperm Capacitation and Acrosome Reaction in Mammalian Sperm. Adv Anat Embryol Cell Biol 220, 93–106. 10.1007/978-3-319-30567-7_5.

16. Doll, S., and Burlingame, A.L. (2015). Mass spectrometry-based detection and assignment of protein posttranslational modifications. ACS Chem Biol 10, 63–71. 10.1021/cb500904b.

17. Riley, N.M., and Coon, J.J. (2016). Phosphoproteomics in the Age of Rapid and Deep Proteome Profiling. Anal Chem 88, 74–94. 10.1021/acs.analchem.5b04123.

18. Ferguson, F.M., and Gray, N.S. (2018). Kinase inhibitors: the road ahead. Nat Rev Drug Discov 17, 353–377. 10.1038/nrd.2018.21.

19. Skerrett-Byrne, D.A., Anderson, A.L., Bromfield, E.G., Bernstein, I.R., Mulhall, J.E., Schjenken, J.E., Dun, M.D., Humphrey, S.J., and Nixon, B. (2022). Global profiling of the proteomic changes associated with the post-testicular maturation of mouse spermatozoa. Cell Rep 41, 111655. 10.1016/j.celrep.2022.111655.

20. Humphrey, S.J., Karayel, O., James, D.E., and Mann, M. (2018). High-throughput and high-sensitivity phosphoproteomics with the EasyPhos platform. Nat Protoc 13, 1897–1916. 10.1038/s41596-018-0014-9.

21. Olsen, J.V., Blagoev, B., Gnad, F., Macek, B., Kumar, C., Mortensen, P., and Mann, M. (2006). Global, in vivo, and site-specific phosphorylation dynamics in signaling networks. Cell 127, 635–648. 10.1016/j.cell.2006.09.026.

22. Consortium, T.U. (2022). UniProt: the Universal Protein Knowledgebase in 2023. Nucleic acids research 51, D523–D531. 10.1093/nar/gkac1052.

23. Skerrett-Byrne, D.A., Pepin, A.S., Laurent, K., Beckers, J., Schneider, R., Hrabě de Angelis, M., and Teperino, R. (2025). Dad’s Diet Shapes the Future: How Paternal Nutrition Impacts Placental Development and Childhood Metabolic Health. Mol Nutr Food Res, e70261. 10.1002/mnfr.70261.

24. Redgrove, K.A., Nixon, B., Baker, M.A., Hetherington, L., Baker, G., Liu, D.-Y., and Aitken, R.J. (2012). The molecular chaperone HSPA2 plays a key role in regulating the expression of sperm surface receptors that mediate sperm-egg recognition. PLoS One 7, e50851.

25. Bromfield, E., Aitken, R.J., and Nixon, B. (2015). Novel characterization of the HSPA2-stabilizing protein BAG6 in human spermatozoa. MHR: Basic science of reproductive medicine 21, 755–769.

26. Skerrett-Byrne, D.A., Stanger, S.J., Trigg, N.A., Anderson, A.L., Sipilä, P., Bernstein, I.R., Lord, T., Schjenken, J.E., Murray, H.C., Verrills, N.M., et al. (2024). Phosphoproteomic analysis of the adaption of epididymal epithelial cells to corticosterone challenge. Andrology 12, 1038–1057. 10.1111/andr.13636.

27. Hornbeck, P.V., Zhang, B., Murray, B., Kornhauser, J.M., Latham, V., and Skrzypek, E. (2015). PhosphoSitePlus, 2014: mutations, PTMs and recalibrations. Nucleic acids research 43, D512–520. 10.1093/nar/gku1267.

28. Hagn, M., Marschall, S., and Hrabè de Angelis, M. (2007). EMMA—The European mouse mutant archive. Briefings in Functional Genomics 6, 186–192. 10.1093/bfgp/elm018.

29. Zhang, X., Tu, H., Zhou, X., Wang, B., Guo, Y., Situ, C., Qi, Y., Li, Y., and Guo, X. (2024). Quantitative Phosphoproteomic Profiling of Mouse Sperm Maturation in Epididymis Revealed Kinases Important for Sperm Motility. Molecular & Cellular Proteomics 23, 100810.

30. Skerget, S., Rosenow, M.A., Petritis, K., and Karr, T.L. (2015). Sperm Proteome Maturation in the Mouse Epididymis. PLoS One 10, e0140650. 10.1371/journal.pone.0140650.

31. Pini, T., Nixon, B., Karr, T.L., Teperino, R., Sanz-Moreno, A., da Silva-Buttkus, P., Tüttelmann, F., Kliesch, S., Gailus-Durner, V., Fuchs, H., et al. (2025). Towards a Kingdom of Reproductive Life – the Core Sperm Proteome. bioRxiv, 2025.2003.2006.641666. 10.1101/2025.03.06.641666.

32. Jumeau, F., Com, E., Lane, L., Duek, P., Lagarrigue, M.l., Lavigne, R.g., Guillot, L., Rondel, K., Gateau, A., and Melaine, N. (2015). Human spermatozoa as a model for detecting missing proteins in the context of the chromosome-centric human proteome project. Journal of proteome research 14, 3606–3620.

33. Gervasi, M.G., and Visconti, P.E. (2016). Chang’s meaning of capacitation: A molecular perspective. Molecular Reproduction and Development 83, 860–874.

34. Aitken, R.J., and Nixon, B. (2013). Sperm capacitation: a distant landscape glimpsed but unexplored. Molecular human reproduction 19, 785–793.

35. Baker, M.A., Smith, N.D., Hetherington, L., Pelzing, M., Condina, M.R., and Aitken, R.J. (2011). Use of titanium dioxide to find phosphopeptide and total protein changes during epididymal sperm maturation. J Proteome Res 10, 1004–1017. 10.1021/pr1007224.

36. Baker, M.A., Hetherington, L., Weinberg, A., Naumovski, N., Velkov, T., Pelzing, M., Dolman, S., Condina, M.R., and Aitken, R.J. (2012). Analysis of phosphopeptide changes as spermatozoa acquire functional competence in the epididymis demonstrates changes in the post-translational modification of Izumo1. J Proteome Res 11, 5252–5264. 10.1021/pr300468m.

37. Humphrey, S.J., Karayel, O., James, D.E., and Mann, M. (2018). High-throughput and high-sensitivity phosphoproteomics with the EasyPhos platform. Nature protocols 13, 1897–1916.

38. Naz, R.K., and Rajesh, P.B. (2004). Role of tyrosine phosphorylation in sperm capacitation / acrosome reaction. Reprod Biol Endocrinol 2, 75. 10.1186/1477-7827-2-75.

39. Cooper, T.G. (2012). The epididymis, sperm maturation and fertilisation (Springer Science & Business Media).

40. Robaire, B., Hinton, B.T., and Orgebin-Crist, M.-C. (2006). CHAPTER 22 - The Epididymis. In Knobil and Neill’s Physiology of Reproduction (Third Edition), J.D. Neill, ed. (Academic Press), pp. 1071–1148. 10.1016/B978-012515400-0/50027-0.

41. Cornwall, G.A. (2009). New insights into epididymal biology and function. Hum Reprod Update 15, 213–227. 10.1093/humupd/dmn055.

42. Gatti, J.L., Castella, S., Dacheux, F., Ecroyd, H., Métayer, S., Thimon, V., and Dacheux, J.L. (2004). Post-testicular sperm environment and fertility. Animal Reproduction Science 82-83, 321-339. 10.1016/j.anireprosci.2004.05.011.

43. Jones, R., and Murdoch, R. (1996). Regulation of the motility and metabolism of spermatozoa for storage in the epididymis of eutherian and marsupial mammals. Reproduction, Fertility and Development 8, 553–568. 10.1071/RD9960553.

44. Steger, K., Klonisch, T., Gavenis, K., Behr, R., Schaller, V., Drabent, B., Doenecke, D., Nieschlag, E., Bergmann, M., And Weinbauer, G.F. (1999). Round spermatids show normal testis specific hit but reduced cAMP responsive element modulator and transition protein 1 expression in men with round spermatid maturation arrest. Journal of andrology 20, 747–754.

45. Björkgren, I., and Sipilä, P. (2019). The impact of epididymal proteins on sperm function. Reproduction 158, R155–R167.

46. Zhang, X., Xiao, Z., Zhang, J., Xu, C., Liu, S., Cheng, L., Zhou, S., Zhao, S., Zhang, Y., Wu, J., et al. (2022). Differential requirements of IQUB for the assembly of radial spoke 1 and the motility of mouse cilia and flagella. Cell Rep 41, 111683. 10.1016/j.celrep.2022.111683.

47. Skerrett-Byrne, D.A., Teperino, R., and Nixon, B. (2024). ShinySperm: navigating the sperm proteome landscape. Reprod Fertil Dev 36. 10.1071/rd24079.

48. Jin, J., Jin, N., Zheng, H., Ro, S., Tafolla, D., Sanders, K.M., and Yan, W. (2007). Catsper3 and Catsper4 are essential for sperm hyperactivated motility and male fertility in the mouse. Biol Reprod 77, 37–44. 10.1095/biolreprod.107.060186.

49. Battistone, M.A., Da Ros, V.G., Salicioni, A.M., Navarrete, F.A., Krapf, D., Visconti, P.E., and Cuasnicú, P.S. (2013). Functional human sperm capacitation requires both bicarbonate-dependent PKA activation and down-regulation of Ser/Thr phosphatases by Src family kinases. Mol Hum Reprod 19, 570–580. 10.1093/molehr/gat033.

50. Fiedler, S.E., Dudiki, T., Vijayaraghavan, S., and Carr, D.W. (2013). Loss of R2D2 proteins ROPN1 and ROPN1L causes defects in murine sperm motility, phosphorylation, and fibrous sheath integrity. Biol Reprod 88, 41. 10.1095/biolreprod.112.105262.

51. Castaneda, J.M., Miyata, H., Archambeault, D.R., Satouh, Y., Yu, Z., Ikawa, M., and Matzuk, M.M. (2020). Mouse t-complex protein 11 is important for progressive motility in sperm†. Biol Reprod 102, 852–862. 10.1093/biolre/ioz226.

52. Richardson, R.T., Yamasaki, N., and Michael, G. (1994). Sequence of a rabbit sperm zona pellucida binding protein and localization during the acrosome reaction. Developmental Biology 165, 688–701.

53. Dun, M.D., Smith, N.D., Baker, M.A., Lin, M., Aitken, R.J., and Nixon, B. (2011). The chaperonin containing TCP1 complex (CCT/TRiC) is involved in mediating sperm-oocyte interaction. J Biol Chem 286, 36875–36887. 10.1074/jbc.M110.188888.

54. Miyata, H., Oura, S., Morohoshi, A., Shimada, K., Mashiko, D., Oyama, Y., Kaneda, Y., Matsumura, T., Abbasi, F., and Ikawa, M. (2021). SPATA33 localizes calcineurin to the mitochondria and regulates sperm motility in mice. Proceedings of the National Academy of Sciences 118, e2106673118. doi:10.1073/pnas.2106673118.

55. Miyata, H., Satouh, Y., Mashiko, D., Muto, M., Nozawa, K., Shiba, K., Fujihara, Y., Isotani, A., Inaba, K., and Ikawa, M. (2015). Sperm calcineurin inhibition prevents mouse fertility with implications for male contraceptive. Science 350, 442–445. doi:10.1126/science.aad0836.

56. Jones, R.C. (1999). To store or mature spermatozoa? The primary role of the epididymis. Int J Androl 22, 57–67. 10.1046/j.1365-2605.1999.00151.x.

57. Zhou, W., De Iuliis, G.N., Dun, M.D., and Nixon, B. (2018). Characteristics of the epididymal luminal environment responsible for sperm maturation and storage. Frontiers in Endocrinology 9, 59.

58. St-Jean, M., Izard, T., and Sygusch, J. (2007). A hydrophobic pocket in the active site of glycolytic aldolase mediates interactions with Wiskott-Aldrich syndrome protein. Journal of Biological Chemistry 282, 14309–14315.

59. Danshina, P.V., Geyer, C.B., Dai, Q., Goulding, E.H., Willis, W.D., Kitto, G.B., McCarrey, J.R., Eddy, E.M., and O’Brien, D.A. (2010). Phosphoglycerate kinase 2 (PGK2) is essential for sperm function and male fertility in mice. Biol Reprod 82, 136–145. 10.1095/biolreprod.109.079699.

60. Fothergill-Gilmore, L.A., and Watson, H.C. (1989). The phosphoglycerate mutases. Adv Enzymol Relat Areas Mol Biol 62, 227–313. 10.1002/9780470123089.ch6.

61. Rodwell, V.W., Towne, J.C., and Grisolia, S. (1957). The kinetic properties of yeast and muscle phosphoglyceric acid mutase. Journal of Biological Chemistry 228, 875–890.

62. Violante, S., Kyaw, A., Kouatli, L., Paladugu, K., Apostolakis, L., Jenks, M., Johnson, A., Sheldon, R.D., Schilmiller, A.L., Visconti, P.E., et al. (2025). Sperm meet the elevated energy demands to attain fertilization competence by increasing flux through aldolase. bioRxiv, 2025.2004.2009.647926. 10.1101/2025.04.09.647926.

63. Signorini, C., Saso, L., Ghareghomi, S., Telkoparan-Akillilar, P., Collodel, G., and Moretti, E. (2024). Redox Homeostasis and Nrf2-Regulated Mechanisms Are Relevant to Male Infertility. Antioxidants (Basel) 13. 10.3390/antiox13020193.

64. Takahashi, N., Davy, P.M., Gardner, L.H., Mathews, J., Yamazaki, Y., and Allsopp, R.C. (2016). Hypoxia Inducible Factor 1 Alpha Is Expressed in Germ Cells throughout the Murine Life Cycle. PLoS One 11, e0154309. 10.1371/journal.pone.0154309.

65. Bakker, W.J., Harris, I.S., and Mak, T.W. (2007). FOXO3a is activated in response to hypoxic stress and inhibits HIF1-induced apoptosis via regulation of CITED2. Mol Cell 28, 941–953. 10.1016/j.molcel.2007.10.035.

66. Aitken, R.J. (2022). Oxidative stress and reproductive function. Reproduction 164, E5–E8.

67. Mo, L., Wu, J., Liu, L., Liu, H., Ou, C., and He, Y. (2024). HIF 1α Affects Sperm Autophagy and Motility. Andrologia 2024, 4575305.

68. Smyth, S.P., Nixon, B., Skerrett-Byrne, D.A., Burke, N.D., and Bromfield, E.G. (2024). Building an Understanding of Proteostasis in Reproductive Cells: The Impact of Reactive Carbonyl Species on Protein Fate. Antioxid Redox Signal 41, 296–321. 10.1089/ars.2023.0314.

69. Wang, J., Zhou, Q., Ding, J., Yin, T., Ye, P., and Zhang, Y. (2022). The conceivable functions of protein ubiquitination and deubiquitination in reproduction. Frontiers in Physiology 13, 886261.

70. Morales, P., Kong, M., Pizarro, E., and Pasten, C. (2003). Participation of the sperm proteasome in human fertilization. Hum Reprod 18, 1010–1017. 10.1093/humrep/deg111.

71. Lishko, P.V., Kirichok, Y., Ren, D., Navarro, B., Chung, J.-J., and Clapham, D.E. (2012). The control of male fertility by spermatozoan ion channels. Annual review of physiology 74, 453–475.

72. Gong, E.Y., Park, E., Lee, H.J., and Lee, K. (2009). Expression of Atp8b3 in murine testis and its characterization as a testis specific P-type ATPase. Reproduction 137, 345–351. 10.1530/rep-08-0048.

73. Balbach, M., Ghanem, L., Violante, S., Kyaw, A., Romarowski, A., Cross, J.R., Visconti, P.E., Levin, L.R., and Buck, J. (2023). Capacitation induces changes in metabolic pathways supporting motility of epididymal and ejaculated sperm. Frontiers in Cell and Developmental Biology Volume 11 - 2023. 10.3389/fcell.2023.1160154.

74. Puga Molina, L.C., Luque, G.M., Balestrini, P.A., Marín-Briggiler, C.I., Romarowski, A., and Buffone, M.G. (2018). Molecular Basis of Human Sperm Capacitation. Front Cell Dev Biol 6, 72. 10.3389/fcell.2018.00072.

75. Reyes-Miguel, T., Roa-Espitia, A.L., Baltiérrez-Hoyos, R., and Hernández-González, E.O. (2020). CDC42 drives RHOA activity and actin polymerization during capacitation. Reproduction 160, 393–404.

76. Baker, M.A., Hetherington, L., and Aitken, R.J. (2006). Identification of SRC as a key PKA-stimulated tyrosine kinase involved in the capacitation-associated hyperactivation of murine spermatozoa. J Cell Sci 119, 3182–3192. 10.1242/jcs.03055.

77. Mitchell, L.A., Nixon, B., Baker, M.A., and Aitken, R.J. (2008). Investigation of the role of SRC in capacitation-associated tyrosine phosphorylation of human spermatozoa. Mol Hum Reprod 14, 235–243. 10.1093/molehr/gan007.

78. Alvau, A., Battistone, M.A., Gervasi, M.G., Navarrete, F.A., Xu, X., Sánchez-Cárdenas, C., De la Vega-Beltran, J.L., Da Ros, V.G., Greer, P.A., Darszon, A., et al. (2016). The tyrosine kinase FER is responsible for the capacitation-associated increase in tyrosine phosphorylation in murine sperm. Development 143, 2325–2333. 10.1242/dev.136499.

79. Salgado-Lucio, M.L., Ramírez-Ramírez, D., Jorge-Cruz, C.Y., Roa-Espitia, A.L., and Hernández-González, E.O. (2020). FAK regulates actin polymerization during sperm capacitation via the ERK2/GEF-H1/RhoA signaling pathway. J Cell Sci 133. 10.1242/jcs.239186.

80. Eid, S., Turk, S., Volkamer, A., Rippmann, F., and Fulle, S. (2017). KinMap: a web-based tool for interactive navigation through human kinome data. BMC Bioinformatics 18, 16. 10.1186/s12859-016-1433-7.

81. Perez-Riverol, Y., Csordas, A., Bai, J., Bernal-Llinares, M., Hewapathirana, S., Kundu, D.J., Inuganti, A., Griss, J., Mayer, G., Eisenacher, M., et al. (2019). The PRIDE database and related tools and resources in 2019: improving support for quantification data. Nucleic acids research 47, D442–d450. 10.1093/nar/gky1106.

82. Dun, M.D., Anderson, A.L., Bromfield, E.G., Asquith, K.L., Emmett, B., McLaughlin, E.A., Aitken, R.J., and Nixon, B. (2012). Investigation of the expression and functional significance of the novel mouse sperm protein, a disintegrin and metalloprotease with thrombospondin type 1 motifs number 10 (ADAMTS10). Int J Androl 35, 572–589. 10.1111/j.1365-2605.2011.01235.x.

83. Nixon, B., MacIntyre, D.A., Mitchell, L.A., Gibbs, G.M., O’Bryan, M., and Aitken, R.J. (2006). The identification of mouse sperm-surface-associated proteins and characterization of their ability to act as decapacitation factors. Biol Reprod 74, 275–287. 10.1095/biolreprod.105.044644.

84. Nixon, B., Stanger, S.J., Mihalas, B.P., Reilly, J.N., Anderson, A.L., Dun, M.D., Tyagi, S., Holt, J.E., and McLaughlin, E.A. (2015). Next Generation Sequencing Analysis Reveals Segmental Patterns of microRNA Expression in Mouse Epididymal Epithelial Cells. PLoS One 10, e0135605. 10.1371/journal.pone.0135605.

85. Nixon, B., Stanger, S.J., Mihalas, B.P., Reilly, J.N., Anderson, A.L., Tyagi, S., Holt, J.E., and McLaughlin, E.A. (2015). The microRNA signature of mouse spermatozoa is substantially modified during epididymal maturation. Biol Reprod 93, 91. 10.1095/biolreprod.115.132209.

86. Anderson, A.L., Stanger, S.J., Mihalas, B.P., Tyagi, S., Holt, J.E., McLaughlin, E.A., and Nixon, B. (2015). Assessment of microRNA expression in mouse epididymal epithelial cells and spermatozoa by next generation sequencing. Genom Data 6, 208–211. 10.1016/j.gdata.2015.09.012.

87. Skerrett-Byrne, D.A., Trigg, N.A., Bromfield, E.G., Dun, M.D., Bernstein, I.R., Anderson, A.L., Stanger, S.J., MacDougall, L.A., Lord, T., Aitken, R.J., et al. (2021). Proteomic Dissection of the Impact of Environmental Exposures on Mouse Seminal Vesicle Function. Mol Cell Proteomics 20, 100107. 10.1016/j.mcpro.2021.100107.

88. Zhou, W., De Iuliis, G.N., Turner, A.P., Reid, A.T., Anderson, A.L., McCluskey, A., McLaughlin, E.A., and Nixon, B. (2017). Developmental expression of the dynamin family of mechanoenzymes in the mouse epididymis. Biol Reprod 96, 159–173. 10.1095/biolreprod.116.145433.

89. Smyth, S.P., Nixon, B., Anderson, A.L., Murray, H.C., Martin, J.H., MacDougall, L.A., Robertson, S.A., Skerrett-Byrne, D.A., and Schjenken, J.E. (2022). Elucidation of the protein composition of mouse seminal vesicle fluid. Proteomics 22, e2100227. 10.1002/pmic.202100227.

90. Rappsilber, J., Mann, M., and Ishihama, Y. (2007). Protocol for micro-purification, enrichment, pre-fractionation and storage of peptides for proteomics using StageTips. Nat Protoc 2, 1896–1906. 10.1038/nprot.2007.261.

91. Bromfield, E.G., Aitken, R.J., Anderson, A.L., McLaughlin, E.A., and Nixon, B. (2015). The impact of oxidative stress on chaperone-mediated human sperm–egg interaction. Human Reproduction 30, 2597–2613.

92. Cafe, S.L., Nixon, B., Dun, M.D., Roman, S.D., Bernstein, I.R., and Bromfield, E.G. (2020). Oxidative Stress Dysregulates Protein Homeostasis Within the Male Germ Line. Antioxid Redox Signal 32, 487–503. 10.1089/ars.2019.7832.

93. Dickinson, M.E., Flenniken, A.M., Ji, X., Teboul, L., Wong, M.D., White, J.K., Meehan, T.F., Weninger, W.J., Westerberg, H., Adissu, H., et al. (2016). High-throughput discovery of novel developmental phenotypes. Nature 537, 508–514. 10.1038/nature19356.

94. Groza, T., Gomez, F.L., Mashhadi, H.H., Muñoz-Fuentes, V., Gunes, O., Wilson, R., Cacheiro, P., Frost, A., Keskivali-Bond, P., Vardal, B., et al. (2022). The International Mouse Phenotyping Consortium: comprehensive knockout phenotyping underpinning the study of human disease. Nucleic Acids Research 51, D1038–D1045. 10.1093/nar/gkac972.

95. Fuchs, H., Aguilar-Pimentel, J.A., Amarie, O.V., Becker, L., Calzada-Wack, J., Cho, Y.L., Garrett, L., Hölter, S.M., Irmler, M., Kistler, M., et al. (2018). Understanding gene functions and disease mechanisms: Phenotyping pipelines in the German Mouse Clinic. Behav Brain Res 352, 187–196. 10.1016/j.bbr.2017.09.048.

96. Nixon, B., De Iuliis, G.N., Hart, H.M., Zhou, W., Mathe, A., Bernstein, I.R., Anderson, A.L., Stanger, S.J., Skerrett-Byrne, D.A., Jamaluddin, M.F.B., et al. (2019). Proteomic Profiling of Mouse Epididymosomes Reveals their Contributions to Post-testicular Sperm Maturation. Mol Cell Proteomics 18, S91–s108. 10.1074/mcp.RA118.000946.

97. Nixon, B., Johnston, S.D., Skerrett-Byrne, D.A., Anderson, A.L., Stanger, S.J., Bromfield, E.G., Martin, J.H., Hansbro, P.M., and Dun, M.D. (2019). Modification of Crocodile Spermatozoa Refutes the Tenet That Post-testicular Sperm Maturation Is Restricted To Mammals. Molecular & cellular proteomics: MCP 18, S58–s76. 10.1074/mcp.RA118.000904.

98. Skerrett-Byrne, D.A., Anderson, A.L., Hulse, L., Wass, C., Dun, M.D., Bromfield, E.G., De Iuliis, G.N., Pyne, M., Nicolson, V., Johnston, S.D., and Nixon, B. (2021). Proteomic analysis of koala (phascolarctos cinereus) spermatozoa and prostatic bodies. Proteomics 21, e2100067. 10.1002/pmic.202100067.

99. Skerrett-Byrne, D.A., Bromfield, E.G., Murray, H.C., Jamaluddin, M.F.B., Jarnicki, A.G., Fricker, M., Essilfie, A.T., Jones, B., Haw, T.J., Hampsey, D., et al. (2021). Time-resolved proteomic profiling of cigarette smoke-induced experimental chronic obstructive pulmonary disease. Respirology 26, 960–973. 10.1111/resp.14111.

100. Tyanova, S., Temu, T., Sinitcyn, P., Carlson, A., Hein, M.Y., Geiger, T., Mann, M., and Cox, J. (2016). The Perseus computational platform for comprehensive analysis of (prote)omics data. Nature Methods 13, 731–740. 10.1038/nmeth.3901.

101. Degryse, S., de Bock, C.E., Demeyer, S., Govaerts, I., Bornschein, S., Verbeke, D., Jacobs, K., Binos, S., Skerrett-Byrne, D.A., Murray, H.C., et al. (2018). Mutant JAK3 phosphoproteomic profiling predicts synergism between JAK3 inhibitors and MEK/BCL2 inhibitors for the treatment of T-cell acute lymphoblastic leukemia. Leukemia 32, 788–800. 10.1038/leu.2017.276.

102. Murray, H.C., Enjeti, A.K., Kahl, R.G.S., Flanagan, H.M., Sillar, J., Skerrett-Byrne, D.A., Al Mazi, J.G., Au, G.G., de Bock, C.E., Evans, K., et al. (2021). Quantitative phosphoproteomics uncovers synergy between DNA-PK and FLT3 inhibitors in acute myeloid leukaemia. Leukemia 35, 1782–1787. 10.1038/s41375-020-01050-y.

103. Kramer, A., Green, J., Pollard, J., Jr., and Tugendreich, S. (2014). Causal analysis approaches in Ingenuity Pathway Analysis. Bioinformatics (Oxford, England) 30, 523–530. 10.1093/bioinformatics/btt703.

104. Skerrett-Byrne, D.A., Nixon, B., Bromfield, E.G., Breen, J., Trigg, N.A., Stanger, S.J., Bernstein, I.R., Anderson, A.L., Lord, T., Aitken, R.J., et al. (2021). Transcriptomic analysis of the seminal vesicle response to the reproductive toxicant acrylamide. BMC Genomics 22, 728. 10.1186/s12864-021-07951-1.

105. Pini, T., Nixon, B., Karr, T.L., Teperino, R., Sanz-Moreno, A., da Silva-Buttkus, P., Tüttelmann, F., Kliesch, S., Gailus-Durner, V., Fuchs, H., et al. (2025). Towards a kingdom of reproductive life - the core sperm proteome. Reproduction 169. 10.1530/rep-25-0105.

